# An Adaptive Flagellar Photoresponse Determines the Dynamics of Accurate Phototactic Steering in *Chlamydomonas*

**DOI:** 10.1101/254714

**Authors:** Kyriacos C Leptos, Maurizio Chioccioli, Silvano Furlan, Adriana I Pesci, Raymond E Goldstein

## Abstract

Our understanding of phototaxis of biflagellates stems almost exclusively from the model alga *Chlamydomonas reinhardtii*, via studies of its flagella, light-sensor and steering. However, no comprehensive model linking all these aspects of its physiology and behavior has been constructed and tested experimentally. Here, we develop such a mathematical model by coupling an adaptive flagellar photoresponse to rigid-body dynamics tailored to details of flagellar beating, and corroborate it with experimental data – at the flagellar and tactic levels – to explain the accurate phototactic steering of this alga. We experimentally validate the hypothesized adaptive flagellar photoresponse using high spatio-temporal resolution methodology on immobilized cells, and corroborate the predicted reorientation dynamics of phototactic swimmers using 3D-tracking of free-swimming cells. Finally, we reconfirm, both theoretically and experimentally, that the adaptive nature of the response has peak fidelity at a frequency of about 1.6 Hz, corresponding to the rotation frequency of the cell body.

## Introduction

Directional non-image-based phototaxis – the ability to change direction of motion in order to reorient with a light stimulus – abounds in motile eukaryotic microorganisms, unicellular and multi-cellular alike. From photosynthetic algae (*Bendix, 1960)* to early-stage larvae of marine zooplankton (*Thorson, 1964)*, phototaxis is such a crucial behavioral response for the survival of these organisms that one is led to hypothesize that organisms must have evolved navigational strategies to reach their goal in a very efficient manner. Photosynthetic algae need to harvest light energy to support their metabolic activities, whereas animal larvae perform phototaxis so that their upward motion can enhance their dispersal.

One of the most intriguing features of non-image-based phototaxis is the ability to navigate towards (or away from) light without the presence of a central nervous system. One of the essential sensory components for directional phototaxis (also known as vectorial phototaxis), is a specialized sensor. This is possible in zooplanktonic larvae via a single rhabdomeric photoreceptor cell *(Jékely et al., 2008)* or in the case of motile photosynthetic micro-organisms such as volvocalean algae, a “light antenna” (*Foster and Smyth, 1980)*, which was generally thought to co-localize with the cellular structure called the *eyespot*, a carotenoid-rich orange stigma. *Foster and Smyth (1980)* theorized that in order for vectorial phototaxis to work, the light antenna has to have directional detection, i.e. detect light only on one side, and that the layers of carotenoid vesicles would act as an interference reflector. This hypothesis was later verified in algae by experiments of eyespot-less mutants that lacked the carotenoid vesicles, but could nevertheless do only negative phototaxis *(Ueki et al., 2016).* Their experiments concomitantly showed that the algal cell bodies can function as convex lenses with refractive indices greater than that of water. For the sake of completeness, it should be noted that in zooplankton the “shading” role of the carotenoid vesicles is filled by a single shading pigment cell *(Jékely et al., 2008).*

**Figure 1.**
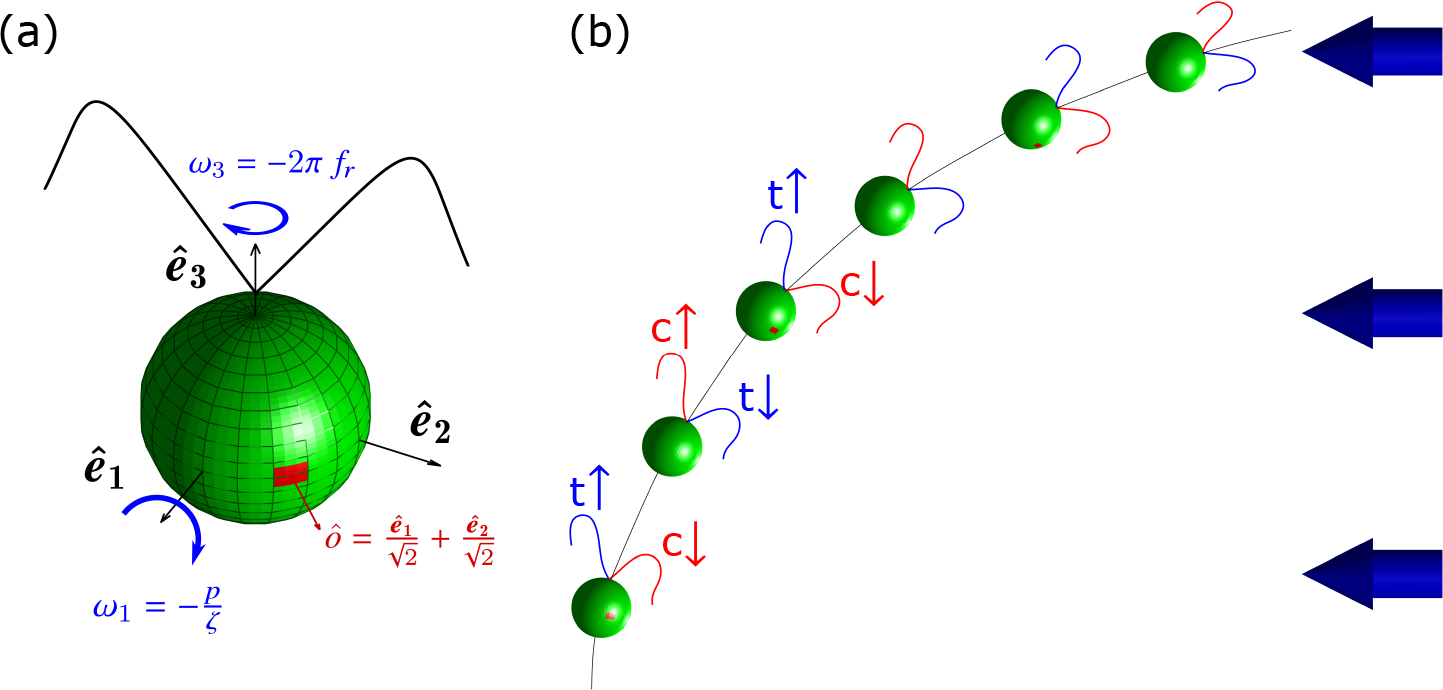
Illustrations of the geometric model of a *Chlamydomonas* cell and of the two-phase model of phototactic activity leading to steering. (a) The axes of the moving frame of the phototactic swimmer is shown, along with the position of the eyespot vector *ô*, shown in red, and found at 45° away from the flagellar beating plane spanned by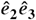. The angular velocities *ω*_1_ and *ω*_3_ are also shown with *p* being the photoresponse, *ζ* a hydrodynamic constant and *f_r_* the frequency of rotation of the cell body, (b) The two phases of photactic activity responsinble for the persistence of phototactic reorientation. t represents the *trans* (in blue) and c the *cis* flagellum (in red).

Among photosynthetic algae the biflagellate species *Chlamydomonas reinhardtii* has been the most studied organism: it exhibits, along with its breast-stroke mode of propagation (*Rüffer and Nultsch, 1985)* and left-handed helix rotation about its axis *(Foster and Smyth, 1980)*, both positive (towards light) and negative (away from light) phototactic responses *(Witman et al., 1993)*, as well as a photoshock/avoidance response. The eyespot in this alga is found on the equator of the cell and at 45° away from the plane of flagellar beating *(Rüffer and Nultsch, 1985).* It was in *Chlamydomonas* that the molecular players mediating phototaxis, the two eyespot-localized photoreceptors, chan-nelrhodopsins A and B, were discovered *(Sineshchekov et al., 2002).* The discovery that these proteins function as light-gated ion channels *(Nagel et al., 2002)*, constituted the initial unraveling of Ariadne’s thread regarding the signal transduction pathway of the photoresponse. Starting instead in the center of this Minoan maze, *Rüffer and Nultsch* used high-speed cinematography to study the flagellar photoresponse *(1990, 1991)*, including the photoshock response *(1995).* With their pioneering work on immobilized *Chlamydomonas* cells they showed, though using a negatively-phototactic strain, that *the front amplitude* of the cells was likely to be responsible for the steering of *Chlamydomonas* towards the light, and that phototaxis is *a result of periodic irradiation and shading.* This result led to the first model for phototaxis *(Schaller et al., 1997)* which divides the turning of the cell into two phases *(Figure 1b): phase I*, in which the rotating eyespot moves from shade to light, causing the flagellum farthest from the eyespot (the *trans* flagellum) to increase its amplitude relative to the flagellum next to the eyespot (the *cis* flagellum), and *phase II*, in which the eyespot moves from light to shade, leading to the two flagella acting in the opposite manner.

Significant contributions to the accurate measuring of flagellar photoresponse at a high temporal resolution were made by *josef etal. (2005)*, who introduced a quadrature photodiode array, a device whose analog signal could be digitized at up to 4000 samples per second. Moreover, this automated method could capture longer time series than previous methods. Despite the limitations of this technology to capture the flagellar photoresponse at high spatial resolution, the authors were able to extract important information regarding flagellar beat-frequency and stroke-velocity.

In recent years, two types of models have sought to describe phototaxis: (i) numerical and theoretical models based on hydrodynamics and heuristic ciliary or flagellar response functions for ciliated larvae *(Jékely et al., 2008*) and biflagellate algae (*Bennett and Golestanian, 2015);* (ii) theoretical adaptation-based models for the green alga *Volvox* (*Drescher et al., 2010)*, the multicelllular “relative” of *Chlamydomonas.* In this study, we have developed a comprehensive mathematical adaptation-based model, in the spirit of *Drescher et al. (2010)* and incorporating information from *Schaller et al. (1997)* and *Rüffer and Nultsch (1991)*, coupled to the dynamics of the yaw, pitch, and so roll of a rigid body In order to describe the three-dimensional phototaxis of *Chlamydomonas* cells. Moreover, we have developed new experimental techniques for capturing the flagellar photoresponse of Immobilized cells at high spatio-temporal resolution and to 3D-track the trajectories of free-swimming phototactic cells. Using these techniques we have measured the time scales involved in photoresponse, adaptation and reorientation that theory dictates are necessary for accurate phototaxis.

## Results

### Capturing flagellar photoresponse and phototactic steering

The flagellar photoresponse of *Chlamydomonas reinhardtii* was captured at high spatio-temporal resolution using the experimental setup shown in *Figure 2a.* This setup builds on previous studies *(Polin et al., 2009; Drescher et al., 2010; Leptos et al., 2013)* with the addition of a much smaller optical fiber (ø50 µm-core) to accommodate for the smaller size of a *Chlamydomonas* cell relative to a *Volvox* spheroid.

The experimental setup *(Figure 2b)* used for phototactic steering featured the following modifications relative to its predecessor *(Drescher et al., 2009)* – either engineered in-house or purchased – for ease and reproducibility: First, the sample chamber could be assembled by the user by clamping two acrylic flanges on a square glass tube in a watertight fashion to prevent leaks. The chamber design allowed a more accurate and easy calibration of the field of view and a simpler and better loading system of the sample via two barbed fittings. Furthermore, the new design of the chamber minimized sample contaminations during experiments. Second, the two 5-Megapixel cameras coupled to objectives with higher total magnification (×16) and larger working distance at the same magnification (48 mm vs. 38 mm at ×2) were used to enhance the image performance.

### Flagellar photoresponse is adaptive

We start by applying a step-up light stimulus. The ability to record the flagellar dynamics of *Chlamydomonas* cells, during light stimulation and at high spatio-temporal resolution, revealed many interesting and important features of the flagellar photoresponse upon a stimulus of this form. Firstly, it corrobrated the fact that change in the waveform of the two flagella was in agreement with previous studies of high-speed cinematography (*Rüffer and Nultsch, 1991)*, i.e. during a step-up response the front amplitude of the *trans* flagellum increases whereas the one of the *cis* flagellum decreases *(Figure 3a-b).* Secondly, it showed that the flagellar photoresponse is adaptive in nature *(Figure 3c and Figure 3–Figure Supplement 2).* For that reason we have employed a mathematical model, previously used to describe adaptive photoresponse in *Volvox (Drescher et al., 2010)*, that relates the adaptive photoresponse *p* to a hidden slow-decaying variable *h* by means of the ordinary differential equations (ODEs):

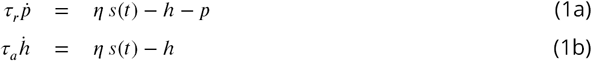

where *s(t)* is the photostimulus function and *η* is a factor with units reciprocal to *s(t).* The hidden variable *h* reflects the internal biochemistry of the cell and is associated with a slower time scale *τ*_0_

**Figure 2.**
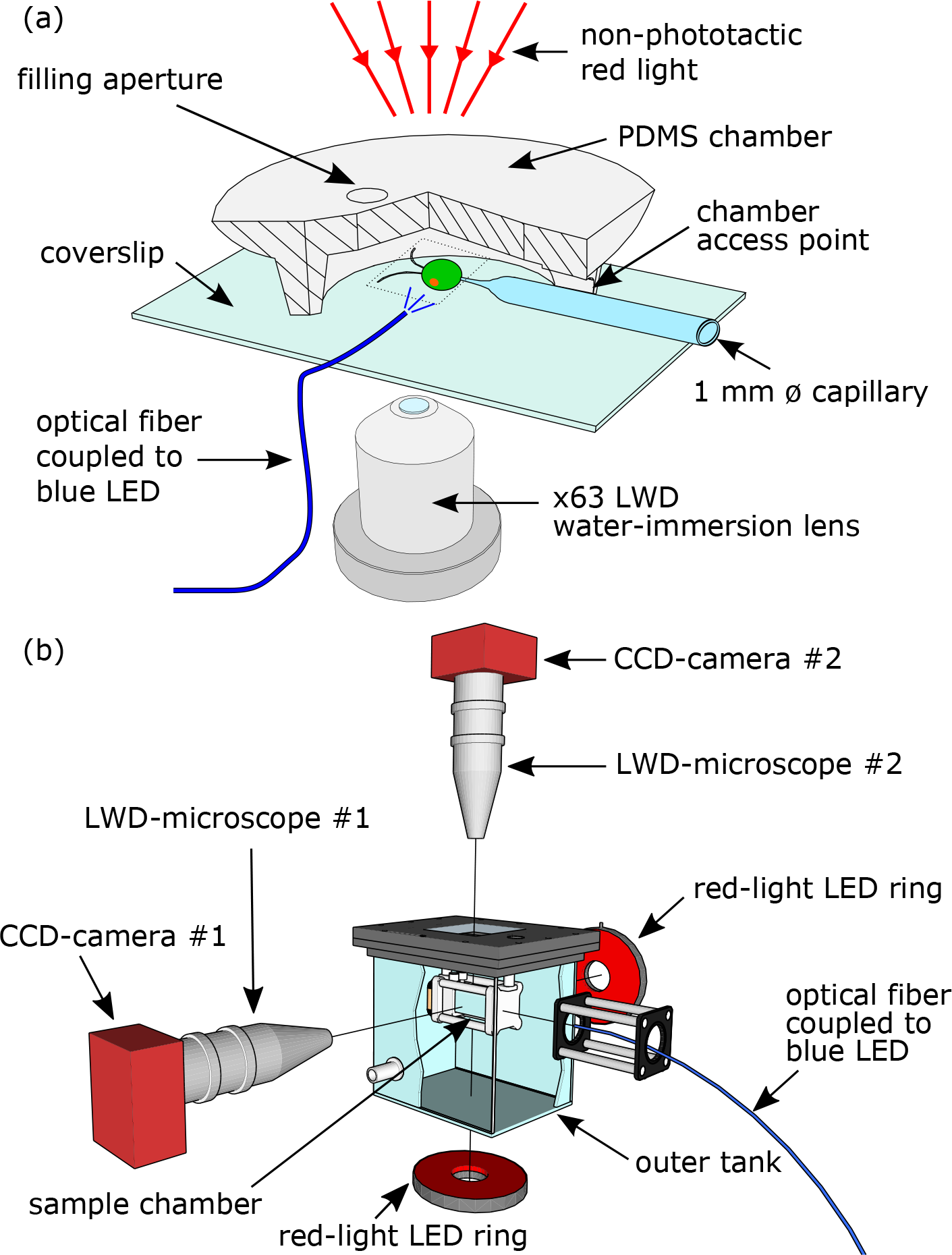
Experimental setups. (a) Experimental setup for measuring the flagellar photoresponse on immobilized cells, inside a PDMS chamber, using a micropipette pulled to a ø5-μm tip. In order to visualize the cell’s beating flagella far from the coverslip, a ×63 LWD objective lens was used. The blue LED used for light stimulation was coupled to a ø5-μm optical fiber, (b) Experimental setup for 3D-tracking phototactic free-swimming cells in a sample chamber immersed into an outer water tank for minimizing thermal convection. Imaging was performed using two aligned LWD microscopes, attached to two CCD cameras.

**Figure 3.**
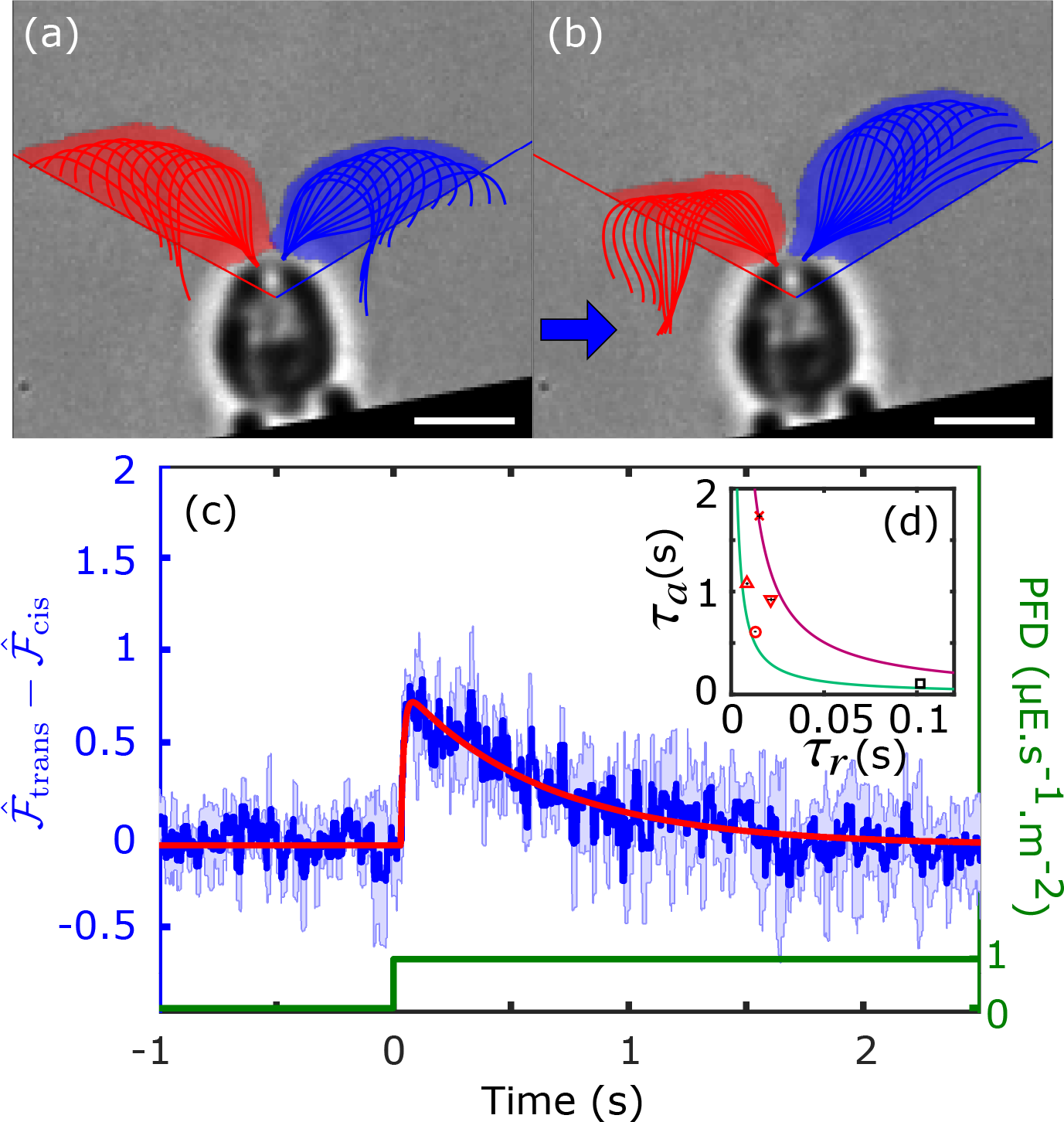
Flagellar photoresponse of immobilized cells upon step-up light stimulation. The raw front amplitudes 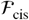 and 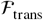 (shaded areas in red for *cis* and blue for *trans*) as defined in the text for each flagellum are shown (a) before and (b) right after the beginning of the photostimulus. The 60° reference lines are also shown. Direction of light is from the left (blue arrow). Scale bar is 5 µm. (c) The mean (dark blue line) and standard deviation (light-blue area) of photoresponse (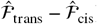) during a step-up stimulus for one cell (*n*_tech_ = 4) fitted to *Equation 2* (red line). (d) Inset showing the mean (red markers) and standard deviation (black error bars) of fitted (*τ_r_, τ_a_*) pairs for *n*_cells_ = 4 upon step-up stimulation. The *(τ_r_, τ_a_)* pair indicated with a black marker is derived from fitting the gain of the frequency response shown in *Figure 4a.* The hyperbolas for 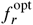 = 1 Hz (red) and 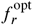 = 2 Hz (green) are also shown.

**Figure 3-Figure supplement 1.**
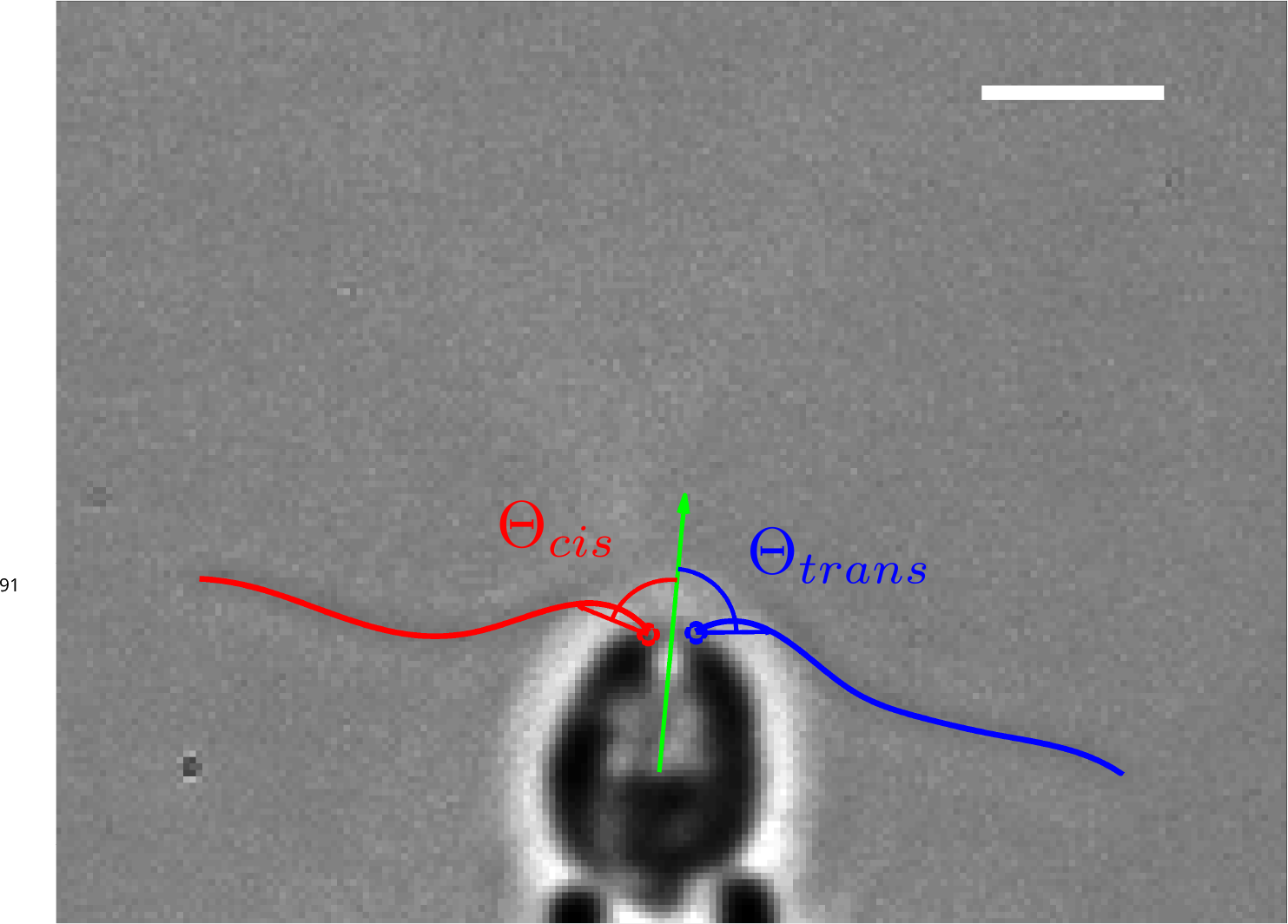
Angle used to define the beginning and the end of a beat. A chord is drawn from the base of each flagellum to a point of fixed length on the flagellum. The angles Θ_cis_ and Θ_trans_ between each of the chords (red for *cis* and blue for *trans* respectively) and the axis of symmetry of the cell (green), were used to define the duration of the flagellar beats. Scale bar is 5µm.

**Figure 3-Figure supplement 2.**
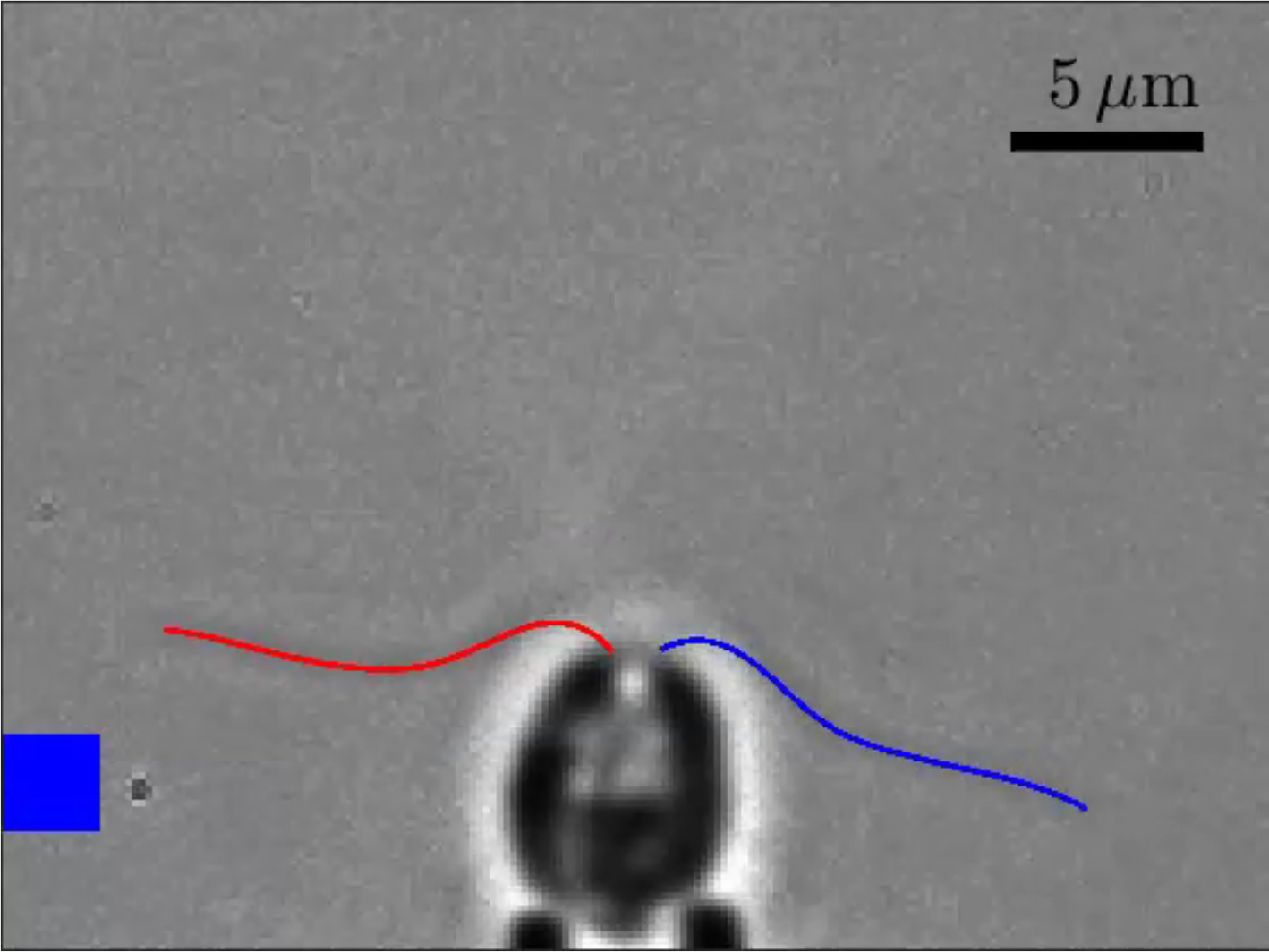
Video showing flagellar photoresponse of immobilized cells upon step-up light stimulation. The optical fiber is illustrated as a grey square that turns blue when stimulus light is turned on. The curves fitted to the *cis* and *trans* flagella are shown in red and blue respectively.

**Figure 3-Figure supplement 3.**
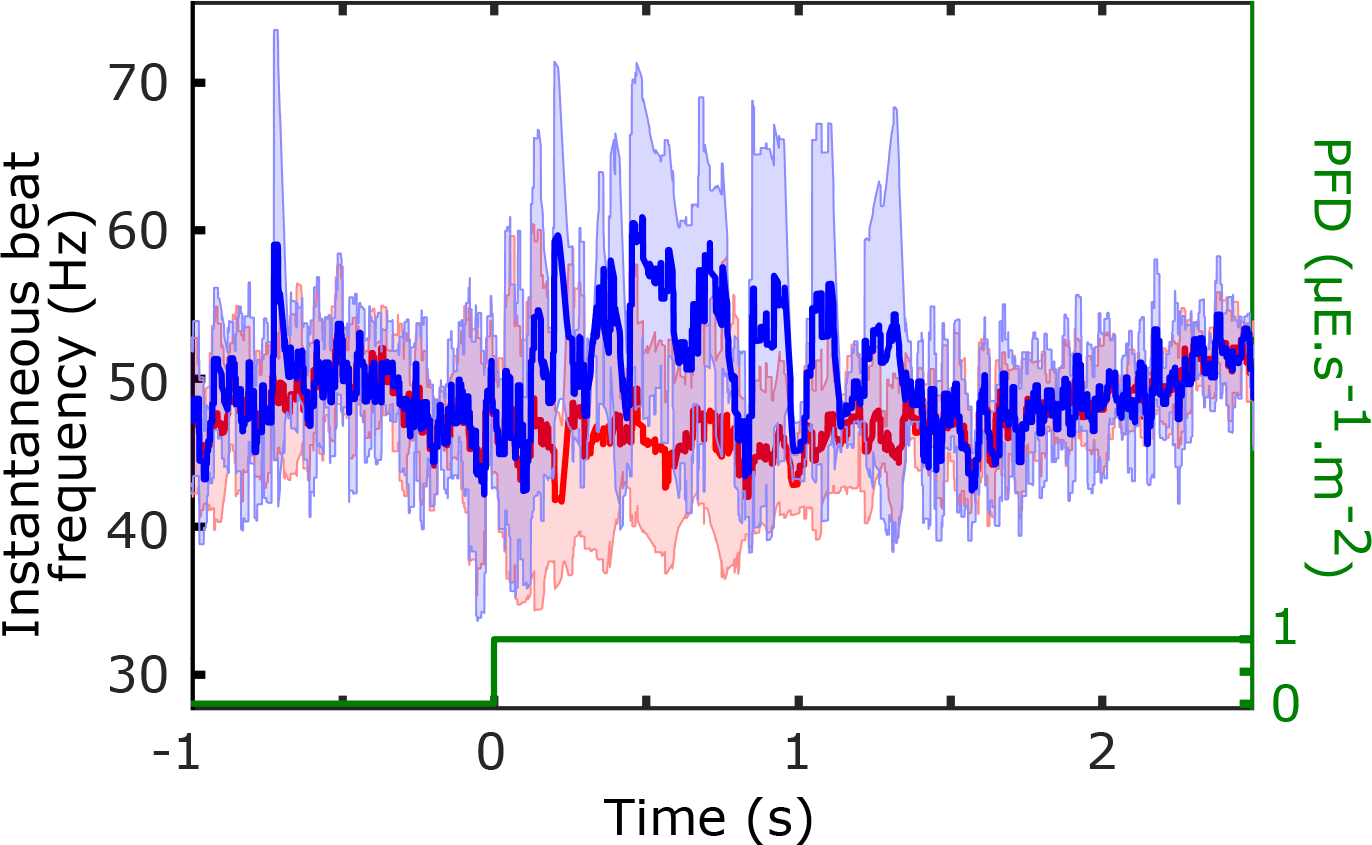
Beat frequency flagellar photoresponse. The beat frequency response for the same cell as shown in Figure 3c averaged over *n_tech_ =* 4 movies. The instantaneous beat frequency was calculated for each beat, ignoring beats that were out of synchrony. The mean and standard deviations of the instantaneous frequencies of the *cis* and *trans* flagella are shown in red and blue respectively.

*(Equation 1b)* than the fast response time scale *τ_r_* (*Equation 1a).* For a step-up stimulus *s(t) = s_0_H(t)*, where *H(t)* is the Heaviside function and *s_0_* is the intensity (flux density) of the light stimulus, these equations can be solved in closed form. Furthermore the data revealed a time-delay of photoresponse upon light stimulation, we therefore add a time delay *t_d_* to this solution to obtain:

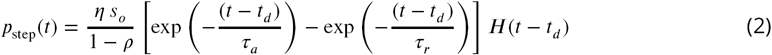

where *ρ = τ_r_/τ_a_.* Experiments clearly show that *ρ <* 1.

We fit the photoresponse data to *Equation 2* to the dimensionless observable 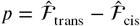, the difference of the *normalized* front amplitudes. The average front amplitude 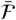 corresponding to each unstimulated cell – used for normalization – was found to be in the range of 35-45 µm^2^. The result of the fittings allowed us to extract values for (*τ_r_, τ_a_*) pairs, as shown in *Figure 3d*, with high accuracy. In order to quantify how these values can affect the efficiency of the photoresponse of a free swimming cell rotating around its central axis, we derived a mathematical relationship relating *τ_r_* and *τ_a_* to the frequency of an oscillating light stimulus *f_s_* (see next section):

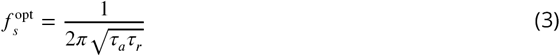

Mathematically this corresponds to the value of *f_s_(= ω_s_/*2*π*), where the gain of rotational frequency response *ℛ(ω_s_)* (described in *Equation 5)* is at its maximum. The relation in *Equation 3* describes a curve (a hyperbola) of optimal (*τ_r_, τ_a_)* pairs for a given stimulus frequency *f_s_* for an immobilized cell, which can be considered equivalents a rotational frequency *f_r_* of a free-swimming cell. As we see from *Figure 3d*, the mean values of fitted (*τ_r_, τ_a_*) pairs along with their standard deviations, for the four cells analyzed, lie within the hyperbolas for 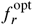 = 1 Hz (red) and 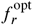 = 2 Hz (green).

Another important feature of the step-up flagellar photoresponse is the time delay between stimulus and response. As shown in *Figure 1a* the eyespot (represented by a red square on the green sphere) is located at an angle *φ* = 45° away from the plane of flagellar beating (located in the 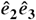 plane). This means that upon light stimulus the flagella of the cell need to be pointing to the same direction as the eyespot for phototaxis to take place in an efficient manner. For that reason we hypothesize that the flagellar response of the cell has been fine-tuned by natural selection to have a delay such that the maximum photoresponse (*p_max_*) will occur after the cell has rotated by an angle of ≈ 45° during its left-handed helix motion. According to *Equation 2* the time *t_max_* at which *p_max_* occurs is:

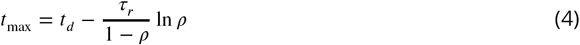

Based on our data we compute *t_max_* to be 94 ± 24 ms (n = 4), which corresponds to an *f_r_ =* 1.1-1.8Hz, assuming a constant *φ* = 45°, or to *φ* = 38° – 64°, assuming a constant 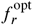 = 1.5 Hz. The range of values for 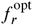 are consistent with the locations of (*τ_r_, τ_a_*) pairs in *Figure 3d.*

### Flagellar photoresponse is fine-tuned with the frequency of rotation of cell body

Cells were stimulated with oscillating light intensity for five different frequencies. If the adaptive photoresponse model holds true, then there should be a maximum response at a resonant frequency corresponding to the frequency of rotation of the cell *f_r_.* This was shown in the past with different techniques, both at the population level (*Yoshimura and Kamiya, 2001)* by measuring the bulk photoreceptor current, and at the single cell-level by *Josef et al. (2006)*, for negative phototaxis and at low spatial resolution. Here we show that this is true at the single cell level, for positive phototaxis and at high spatial resolution, by directly measuring the flagellar photoresponse *p* as defined in the previous section. The results from individual experiments *(Figure 4b-d)* immediately revealed two major findings: (a) The flagellar photoresponse oscillates with the same frequency as the frequency of the amplitude of the light-stimulus. This means that the response is linear and can be described by the solution *p_oscill_* (*Equation 10* in *Materials and Methods)* of the governing equations (*Equation 1a* and *Equation 1b)*, for an oscillating light stimulus *s(t)* (green line in *Figure 4b-d).* (b)The amplitude of the observed *p* (blue line in *Figure 4b-d)* is higher at certain frequencies than others.

**Figure 4.**
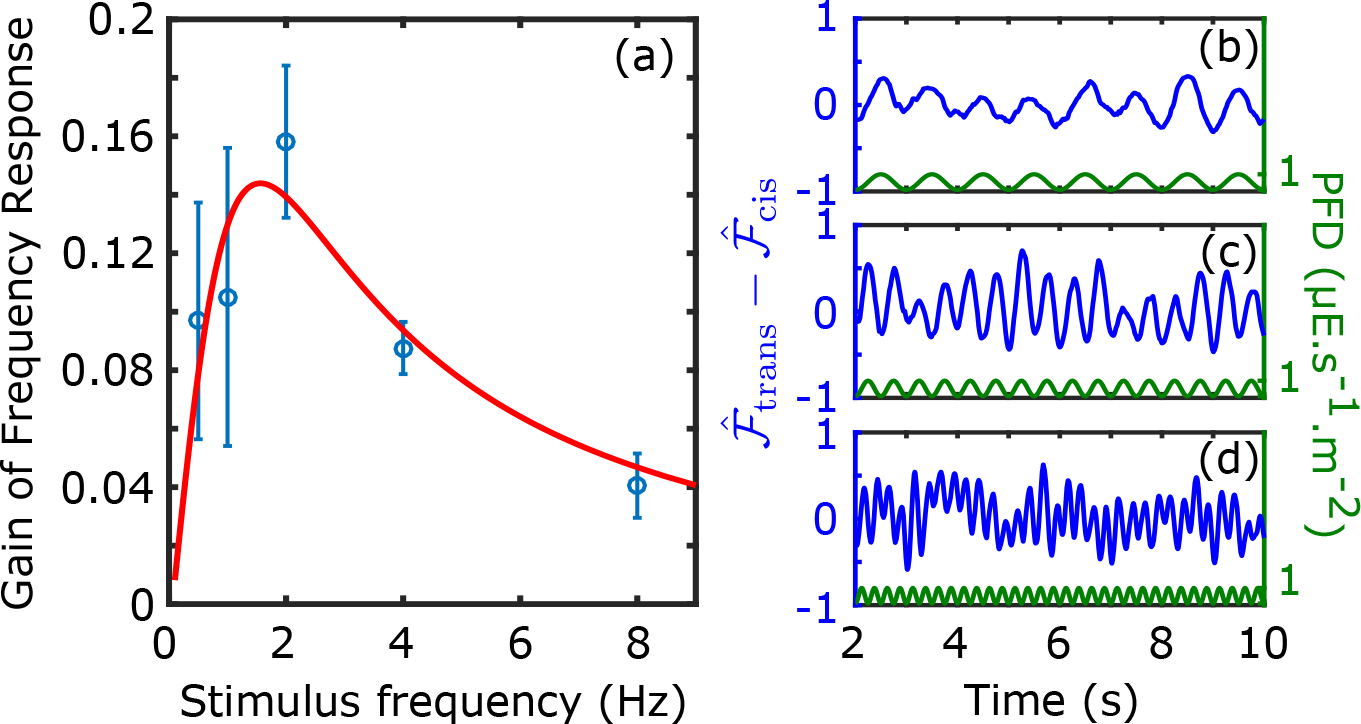
Frequency response of immobilized cells stimulated with oscillating-amplitude light. (a) The calculated gain of the frequency response (for positive phototaxis) for five stimulus frequencies (0.5,1,2,4, and 8 Hz) for *n_cells_* = (blue) fitted to *Equation 5* (red line). The photoresponse (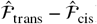) shown (in blue) for three different stimulus frequencies (in green): 1 Hz (b), 2 Hz (c) and 4 Hz (d). The values of *p* for flagellar beats during instantaneous asynchronies were replaced by interpolated values based on neighbouring synchronous beats. The percentage of asynchronous beats during the time intervals shown were 7.6%, 10.6% and 35.3% for (b), (c) and (d) respectively.

In order to investigate which of the five stimulus frequencies *(f_s_)* gives the most prominent flagellar photoreponse *p* we first derived a relationship (*ℛ*) between *f_s_* (= *ω_s_/*2*π*) and the magnitude of the Fourier transform of *p_oscill_*. This is defined in *Equation 11* of the *Materials and Methods.* The result of the computation, which we refer to as the *gain of the frequency response*, is a function of *ω_s_:*

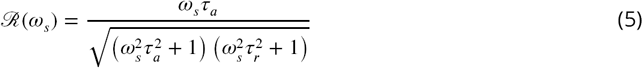

At the experimental level, we calculated the observed gain of the frequency response (blue in *Figure 4a)*, using a Discrete Fourier Transform on the observed *p.* The mean observed gain peaks at 2 Hz. The data were fitted to *Equation 5* giving *τ_r_* ≈ *τ_a_* = 0.1 s, which peaks at ≈1.6 Hz (red in *Figure 4a).*

### Model of phototactic swimmers in three dimensions

Naturally, the information gained from measuring the photoresponse of immobilized cells can be used, initially at least, to get an estimate of the angular velocity *ω_1_* of the cell (*Figure 1a)* during a phototactic turn. In particular, we would like to estimate the angle by which a free-swimming cell – starting at a direction of 90° to the light source – would turn during the first half turn of the rotation of the cell body about 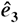 (*Figure 1a).* For pedagogical reasons we provide a more detailed calculation of this estimate in *Appendix 1* as motivation for the full mathematical model that follows.

For this estimate we a consider a simplified swimmer in Stokes flow. The swimmer is composed of a spherical body of radius *R*, and two “flagella” in the shape of thin rods of length *L.* We compute the total torque (*Equation 16* in *Appendix 1)* generated by each of the two flagella – during the effective stroke of the beat – to be *τ*_1,2_ = (2/3)*ζ*_⊥_*f*_*b*_*a*_1,2_*L*^2^, where *f_b_* is the frequency of flagellar beating, *ζ*_⊥_ is the perpendicular viscous drag coefficient, and *a_1,2_* is the amplitude of each flagellum. This expression can also be related to the area swept by each flagellum *A_1,2_* (*Appendix 1-Figure 1)* as *τ_1,2_* = (4/3)*ζ_⊥1_f_b_A_1,2_L*. The physical quantity that causes the cell to turn with angular velocity *ω_1_* about 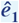 is the difference In the torques (Δ*τ* = *τ*_1_ – *τ*_2_) generated by the two flagella, divided by the rotational drag coefficient *ζ_r_.* Assuming that during a photoresponse the amplitude of flagellum 1 (*a*_1_) is equal to *a + b* and of flagellum 2 (*a*_2_) is equal to *a – b*, where *b* is the amplitude difference from the unstimulated state *a*, then *ω_1_* = -(*τ/ζ_r_*)(2*b/a*). We know that the flagellar amplitude oscillates if it experiences an oscillating stimulus *(Figure 4c)* so if *b(t)* = 2*b_o_* sin (2*πf_r_t)*, where 2*b_o_* is the maximum flagellar amplitude difference and *f_r_* is the frequency of rotation of the body of the cell, then

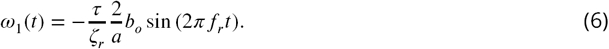

If we integrate Equation 6 for the first half turn (HT) we will obtain the angle of phototactic turning (Φ_HT_) for that period of time during which *b_0_* is assumed to be constant. The result of the integration (Appendix 1) is

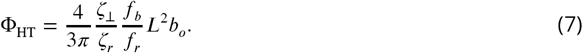

If we substitute *L* = 10 µm, *b_0_* = 1 µm, *a* = 5 µm, ζ_⊥_ = 2.6 × 10^-3^ Pa·s *(Appendix 1), τ_r_* = 3 pN·µm·s *(Appendix 1), f_b_* = 50 Hz and *f_r_* = 2Hz into *Equation 7* we find Φ_HT_ ≈ 0.9rad ≈ 52°. This means that even with this oversimplified model of rod-shaped flagella, where the torque generated is overestimated, it is possible for the phototactic swimmer to reorient with the light source (i.e. turn 52° about 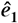) within half turn of cell-body rotation (I.e. a turn of 180° about 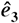). Even though we do not formally define the dimensionless variable *p* – used previously to describe the photoresponse – in terms of the torques (*τ_1,2_*) generated by the two flagella, we nevertheless proceed to utilize this convenient variable to model the reorientation of the phototactic swimmers In three dimensions, by defining *ω_1_* to be proportional to *p.*

The reorientation of phototactic swimmers – in three dimensions – can be described as a system of five nonlinear ODEs expressing, in addition to that of *p* and *h (Equation 1a* and *Equation 1b)*, the time evolution of the three Euler angles of precession (*ϕ*), nutation *(θ)* and rotation *(ψ) (Symon, 1971).* This is achieved by coupling the light stimulus *s(t)* with the amount of light received by the eyespot as the cell turns and rotates. Moreover, the coupling of the Euler angle dynamics to the photoresponse is achieved with the relation *ω_1_ = -(1/ζ)p*, where *ζ* is an effective viscosity, as shown In *Figure 1a.* We postponed the detailed derivation to *Appendix 2.*

Using the assumption that the swimmer’s U-turn lies in a plane *(Appendix 2)*, we reduce the problem to a system of three ODEs in which *ϕ* + *π/2* describes the angle between the direction of the swimmer and the direction of the light stimulus. Moreover, we non-dimensionalize the equations by rescaling time to 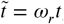 where *ω_r_ =* 2*πf_r_ (Appendix2-Equations 19a-c).*

One of the most important features of phototaxis in *Chlamydomonas* is the separation of time scales between the duration of individual flagellar beats (1*/f_b_* ~ 0.02s), the half-period of cell body rotation (1/2*f_r_* ~ 0.25s) and the time for phototactic reorientation (~ 2 s). As It takes many half-periods of body rotation to execute a turn, we can consider the angle *ϕ* that defines the instantaneous angle between the cell body and the light direction to be approximately constant during each half-period. Under this assumption, we may recast the phototactic dynamics as an iterated map *(Figure 5* and *Figure 5–Figure Supplement 1)* for the quantity Φ_*n*_, defined to be the Euler angle *ϕ* at the end of each half-turn.

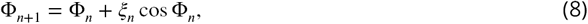

where *ξ_n_* is defined in *Appendix 2-Equation 21* in terms of *n* and the fundamental parameters *(τ_r_, τ_a_, f_r_, ζ, s_0_η).* With the angle *ϕ* as defined above, we see that when a cell swims directly towards the light, Φ = *π/2.* One natural question is whether a cell can reach that orientation from any initial condition in which the eyespot receives some light, corresponding to the initial angle Φ_0_ lying in the range 0 ⩽ Φ_0_ ⩽ *π.* In the usual manner of interpreting such iterated maps, if we choose Φ_0_ = 0, as in *Figure 5*, the angle Φ_1_ after one half-turn is obtained by moving along the vertical blue line from the value Φ_0_ on the horizontal axis until intersecting the green curve Φ_*n*_ + *ξ*_0_ cos Φ_*n*_. Using Φ_1_ so obtained for the next iteration is equivalent to reflecting the blue line off the grey diagonal up to the black curve Φ_*n*_ + *ξ*_1_ cos Φ_*n*_, thus obtaining Φ_2_. Continuing this “cobwebbing”, we see the trajectory converge to Φ = *π/2*, which is a stable fixed point. Choosing Φ_0_ = *π* leads to the cobwebbing trajectory shown in red, which also converges to Φ = *π/2.* This serves as a simple, heuristic demonstration of the manner in which an adaptive photoresponse leads to robust phototaxis.

**Figure 5.**
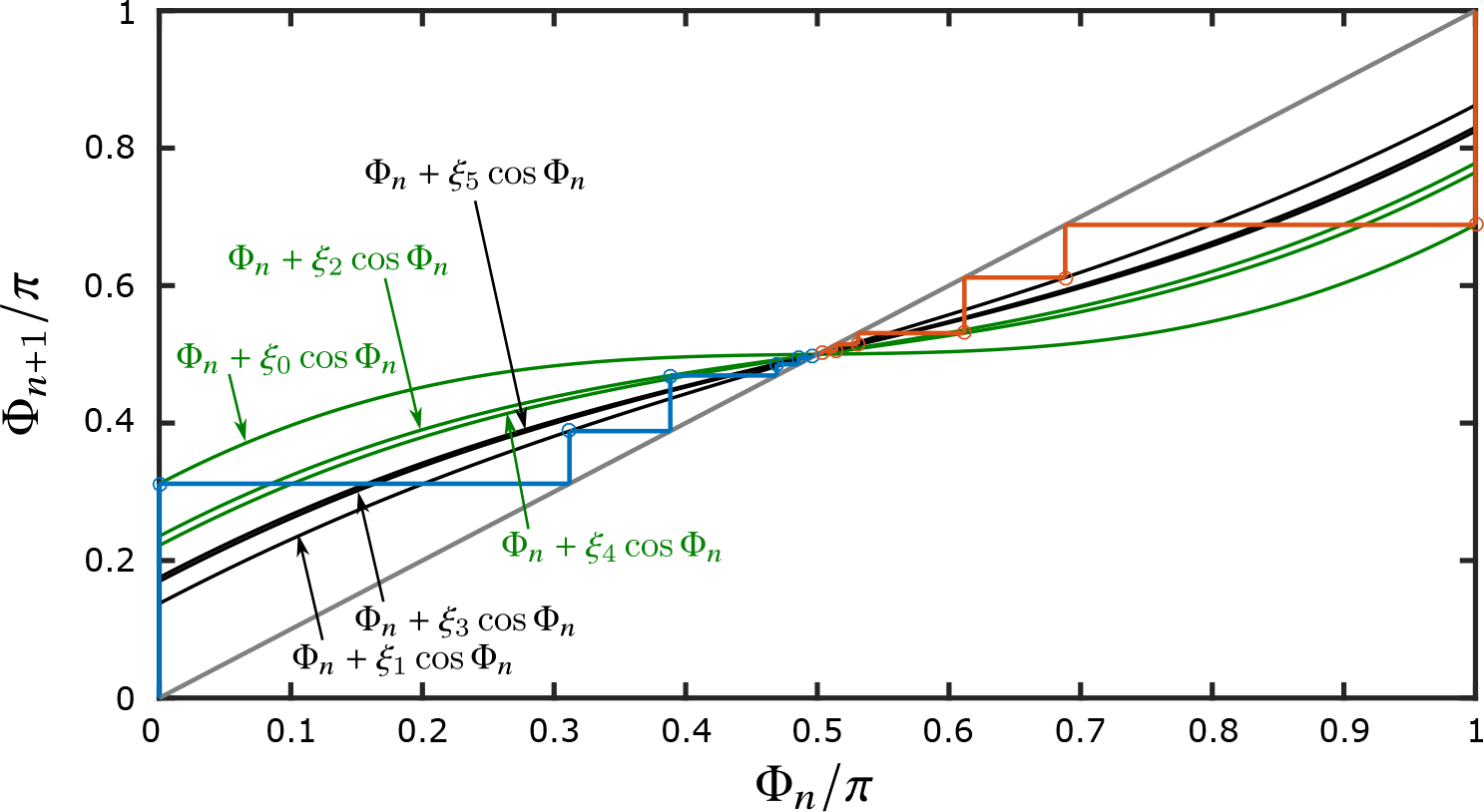
Iterated map of the reorientation model. The iterated map describes the dynamics of two cells, one with initial conditions Φ_0_ = 0 (blue) and the other with initial conditions Φ_0_ = *π* (red), reorienting towards a fixed light source, as described by *Equation 8.* The complete alignment with the light source is described by the fixed point *π/2.* The function Φ_n+1_ = Φ_*n*_ + *ξ_n_* cos Φ_*n*_, is shown for six half turns, i.e. 0 ⩽ *n* ⩽ 5, with even numbers shown in green and odd numbers shown in black. The parameters defining *ξ_n_* (*τ_r_* = 0.06 s, *f_r_* = 1.6 Hz and *σ/ζ* = 10 s^-1^) were chosen to be comparable to the experimental data, while the value *τ_a_* = 0.s was chosen larger than in experiment in order to separate the individual curves to illustrate more clearly the nature of the map.

**Figure 5-Figure supplement 1.**
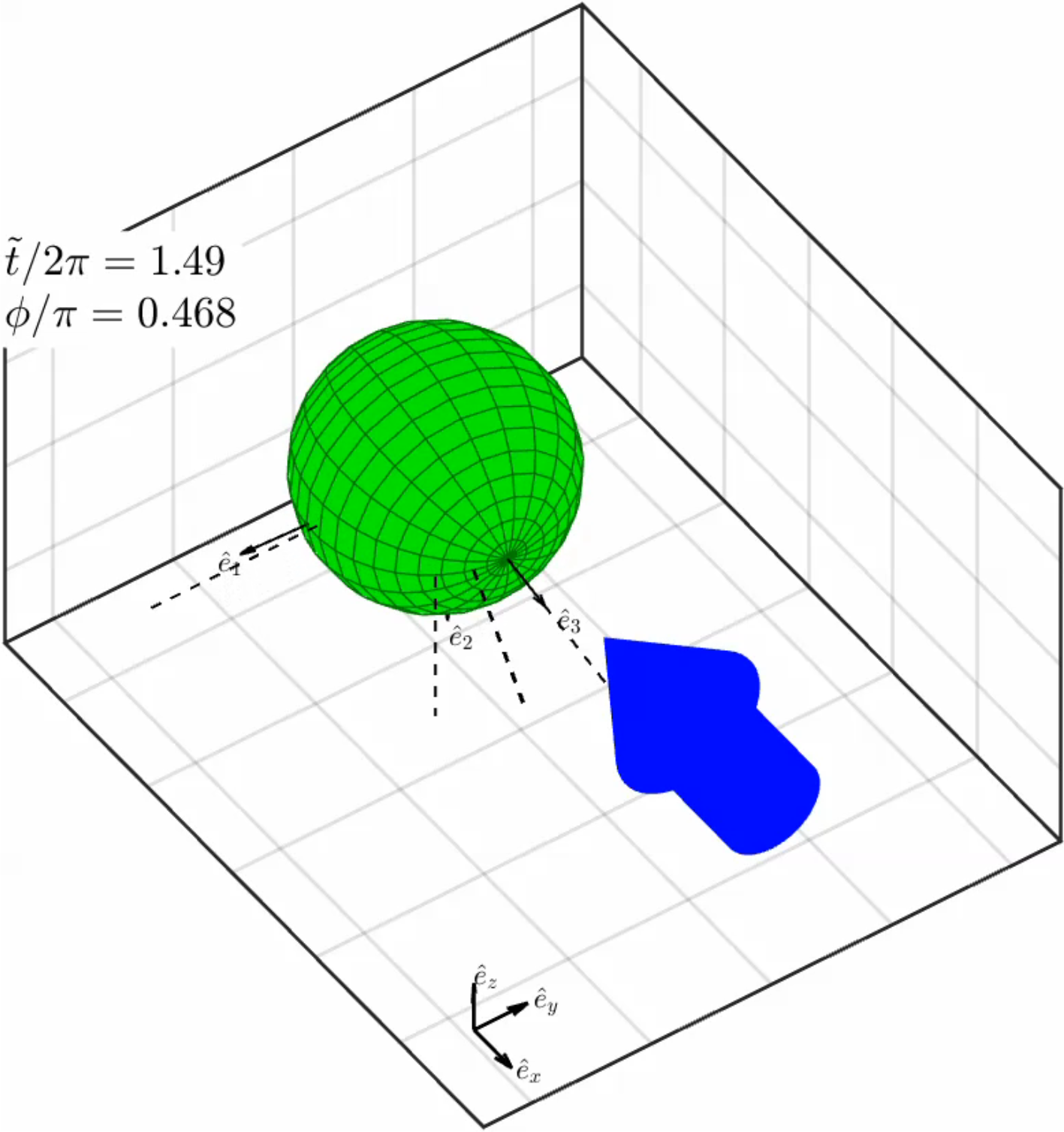
Animation of a solution of an iterated map. Video animation of the reorientation dynamics – to Φ_*n*_ = *π/2* as shown in Figure 5 for the cell with Φ_*0*_ = 0. The position of the vector 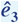 is marked with a dashed line for every half turn. Time is displayed in numbers of full turns (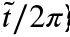) and the interpolated Euler angle *ϕ* is shown in units of *π* radians.

**Figure 5-Figure supplement 2.**
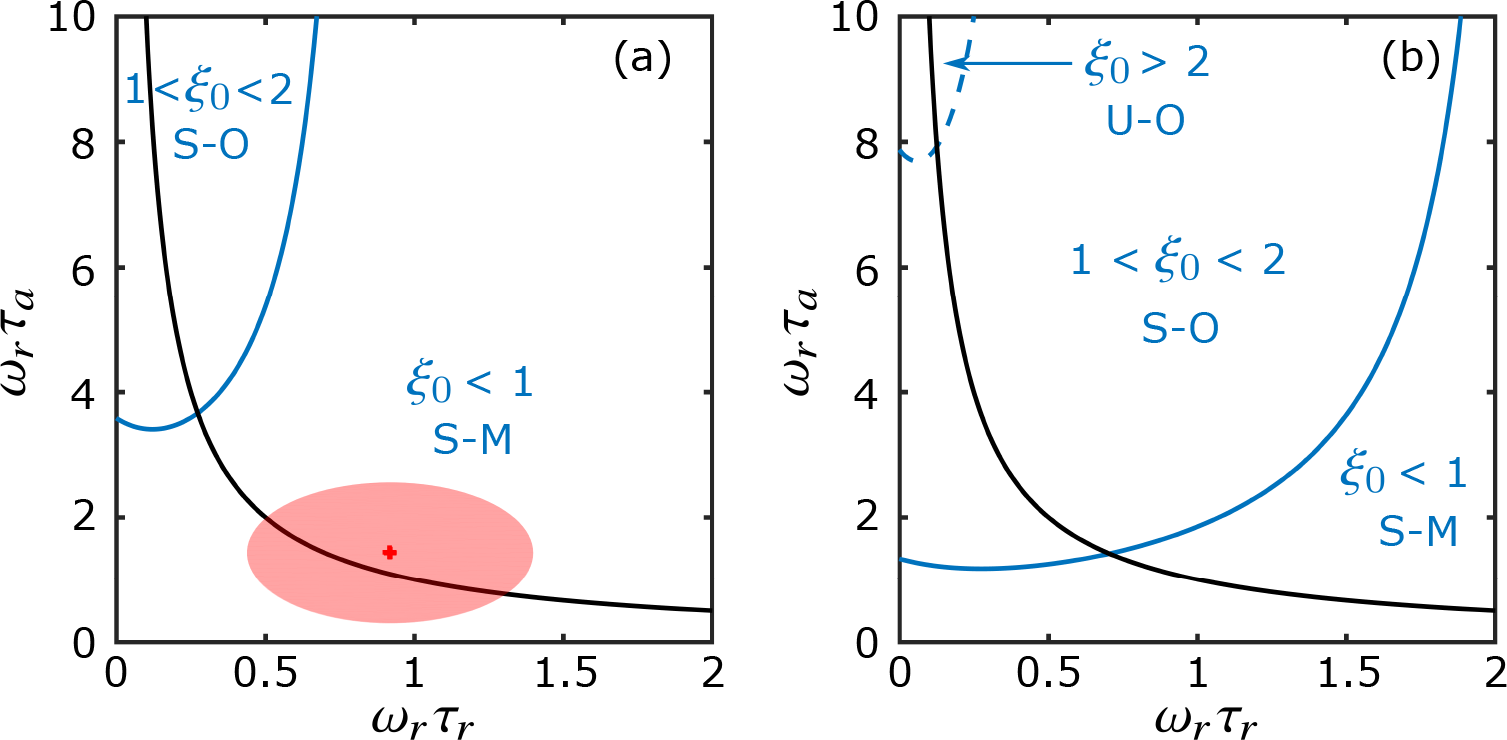
Phase diagram of the dynamics as defined by the values *ξ*_0_. The values of *ξ*_0_, which are plotted as a function of *ω_r_τ_r_* and *ω_r_τ_a_* determine the behavior of the iterated map. Numerically-calculated boundaries where *ξ*_0_ = 1 and *ξ*_0_ = 2 are shown in solid and dashed blue lines respectively. The two phase diagrams presented were generated using two different values of *σ/ζ*: 9 s^-1^ (a) and 15 s^-1^ (b). Regions are labeled as S-M for stable monotonic, S-O for stable oscillatory and U-O for unstable oscillatory. The optimal (rescaled) *τ_r_,τ_a_* pairs – for immobilized cells – are shown as the black line *ω_r_τ_a_ = 1/(ω_r_τ_r_).* Red cross and orange ellipse in (a) summarize the mean and standard deviation of the experimental data on *τ_r_,τ_a_* pairs and *f_r_* shown respectively in Figure 6c and *Figure 6-Figure Supplement 2a.*

**Figure 5-Figure supplement 3.**
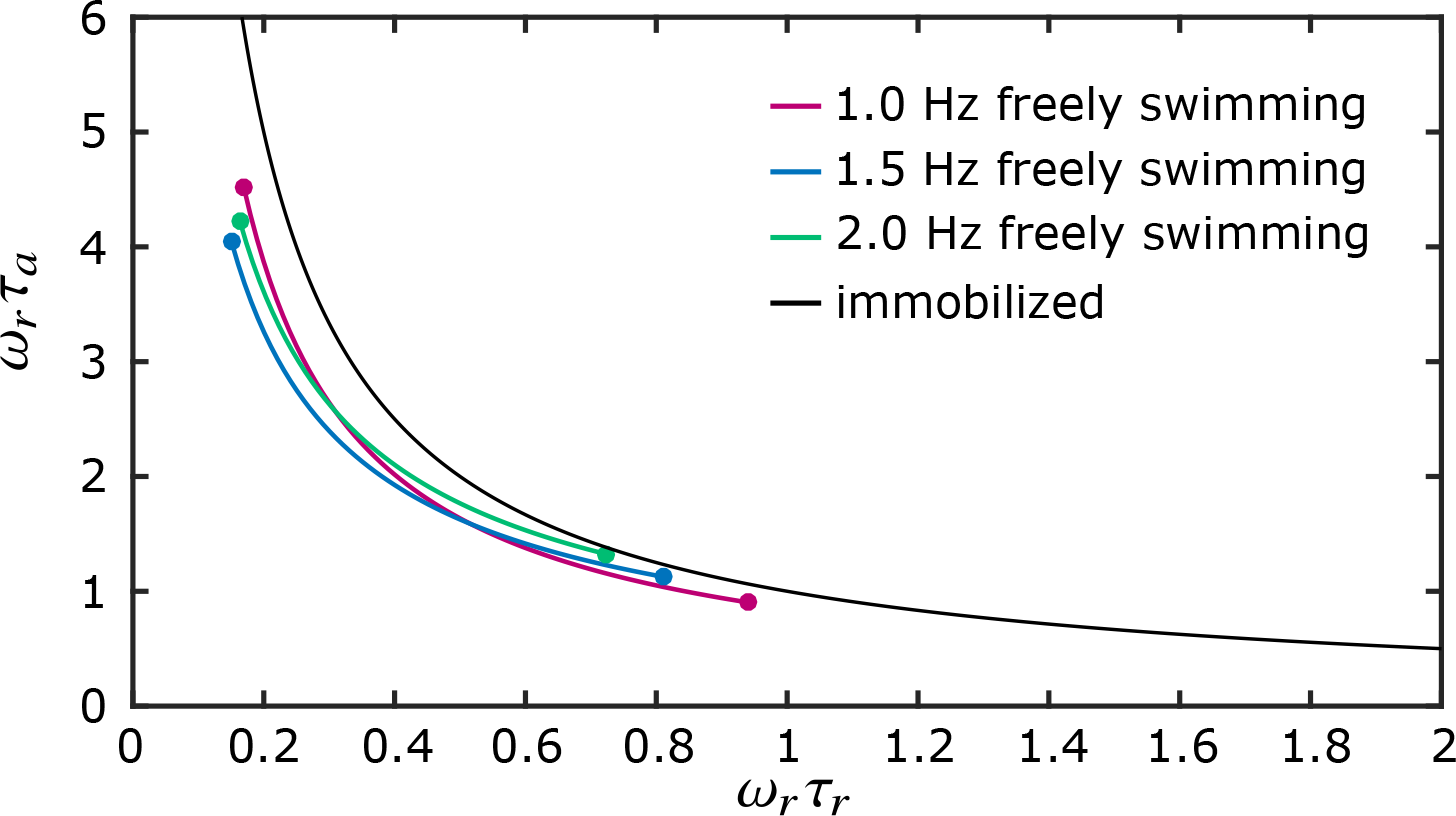
Optimal *τ_r_, τ_a_* pairs extracted from the 3D reorientation model. The optimal (rescaled) *τ_r_,τ_a_* pairs as extracted from the 3D reorientation model when *〈ξ〉 = (ξ_0_ + ξ_1_)/2* is at a maximum for a given *f_r_* and *σ/ζ*. The locus of these points is illustrated with three different fitted line segments which correspond to different values of *f_r_.* The optimal rescaled *τ_r_,τ_a_* pairs – for immobilized cells – are shown as the black line *ω_r_τ_a_ = 1/(ω_r_τ_r_).*

The rate of alignment of the swimming direction of the cell with respect to the light vector can be deduced by setting Φ_n_ = *π/2* – ψ_*n*_, where the angular deviation ψ_*n*_ obeys ψ_*n*+1_ = ψ_*n*_ – *ξ*_*n*_ sin ψ_*n*_ ≃ (1 – *ξ*_*n*_)ψ_*n*_, the latter relation holding for near-alignment (ψ_*n*_ ≪ 1). Heuristically, we can ignore the small differences between the and obtain the approximate solution ψ_*n*_ ~ (1- *ξ*_0_)^n^ψ_0_, which shows that the magnitude of serves as a measure of the rate of reorientation of the cell; If 0 < *ξ*_0_ < 1 the approach is monotonic, if 1 < *ξ*_0_ < 2 it is oscillatory but stable, and if *ξ*_0_ > 2 the reorientation does not occur – the aligned state is unstable (and oscillatory). *Figure 5-Figure Supplement 2* shows these different regimes in the parameters space of *τ_r_* and *τ_a_* for two different values of the prefactor *σ/ζ*, along with the relation *(ω_r_τ_a_)(ω_r_τ_r_)* = 1 of *Equation 3*, which defines the optimum response of immobilized cells. *Figure 5–Figure Supplement 3* shows a comparison between the latter and the results of optimizing the 3D-reorientation rate by maximizing the average 〈*ξ*〉 = (*ξ*_0_ + *ξ*_1_)/and we see the two approaches lead to remarkably similar results.

### Three-dimensional trajectories yield optimized photoresponse parameters

Within the M = 6 pairs of recorded movies, we tracked 283 trajectories with durations greater than 10 s and which included the trigger frame. From those, 44 showed both positive phototaxis and included a full turn to Ω = *π* as shown in *Figure 6a* and *Figure 6–Figure Supplement 1.* These three-dimensional trajectories were cropped to any points for which *π/2* ⩽ Ω ⩽ *π* and which could then be fitted to *Equation 8*, using the relation Ω = Φ + *π/2*, as shown in *Figure 6b.* Out of these, 21 trajectories had good fits (see Methods section for criterion) and the estimated four parameters (*τ_r_, τ_a_, f_r_* = *ω_r_*/2*π* and *μ* = *σ/ζ*) in *ξ_n_* (*Equation 21)* converged to a sufficiently narrow range of values. More specifically, pairs of the means of fitted *τ_r_* and *τ_0_* (*Figure 6c)* fall within the values of optimal response and adaptation time scales as described by *Equation 3*, mostly between the hyperbolas 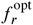 = 1.5 Hz (blue line) and 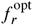 = 2 Hz (green line). The distributions of the means of fitted values for the other two parameters *f_r_* and *μ* are shown in *Figure 6-Figure Supplement 2a* and *Figure 6 Figure Supplement 2d* respectively. The median value for the fitted rotational frequency of the cell (*f*_r_) was found to be 1.78 Hz, in strong agreement with the maximum value of the fitted gain of frequency response in *Figure 4a.* Parameter *μ* had a median value of 8.98 s^-1^ and it was found to be independent for the two different light intensities used. Finally, if we perform a global average of the values for *f_r_, τ_r_*, and *τ_a_* for the data shown in *Figure 6c* we can locate it on the stability diagram in *Figure 5-Figure Supplement 2a*, where we see that *Chlamydomonas* operates very close to the optimum photoresponse curve, well within the stable-monotonic regime of alignment.

## Discussion

This study has achieved three major goals: the development of modern methods to capture flagellar photoresponse at high spatio-temporal resolution, the measurement of important biochemical time scales for the understanding of phototaxis and lastly the integration of the above information through the development of a biochemistry-based model to accurately describe the phototactic behavior of *Chlamydomonas* in terms of the dynamics of reorientation to the light source in three dimensions.

In addition, this study has addressed issues relating to past observations: With respect to the lag time *t_d_* of the photoresponse, we have measured a value of 32 ± 9 ms (n = 4), very similar to the value 30-40 ms observed by *Rüffer and Nultsch (1991).* In addition, we argue that the maximum flagellar response would take place at *t_max_* as shown In *Equation 4*, which adds a correction factor to *t_d_.* This is important when assessing the efficiency of the response with respect to the frequency of rotation of the cell body.

**Figure 6.**
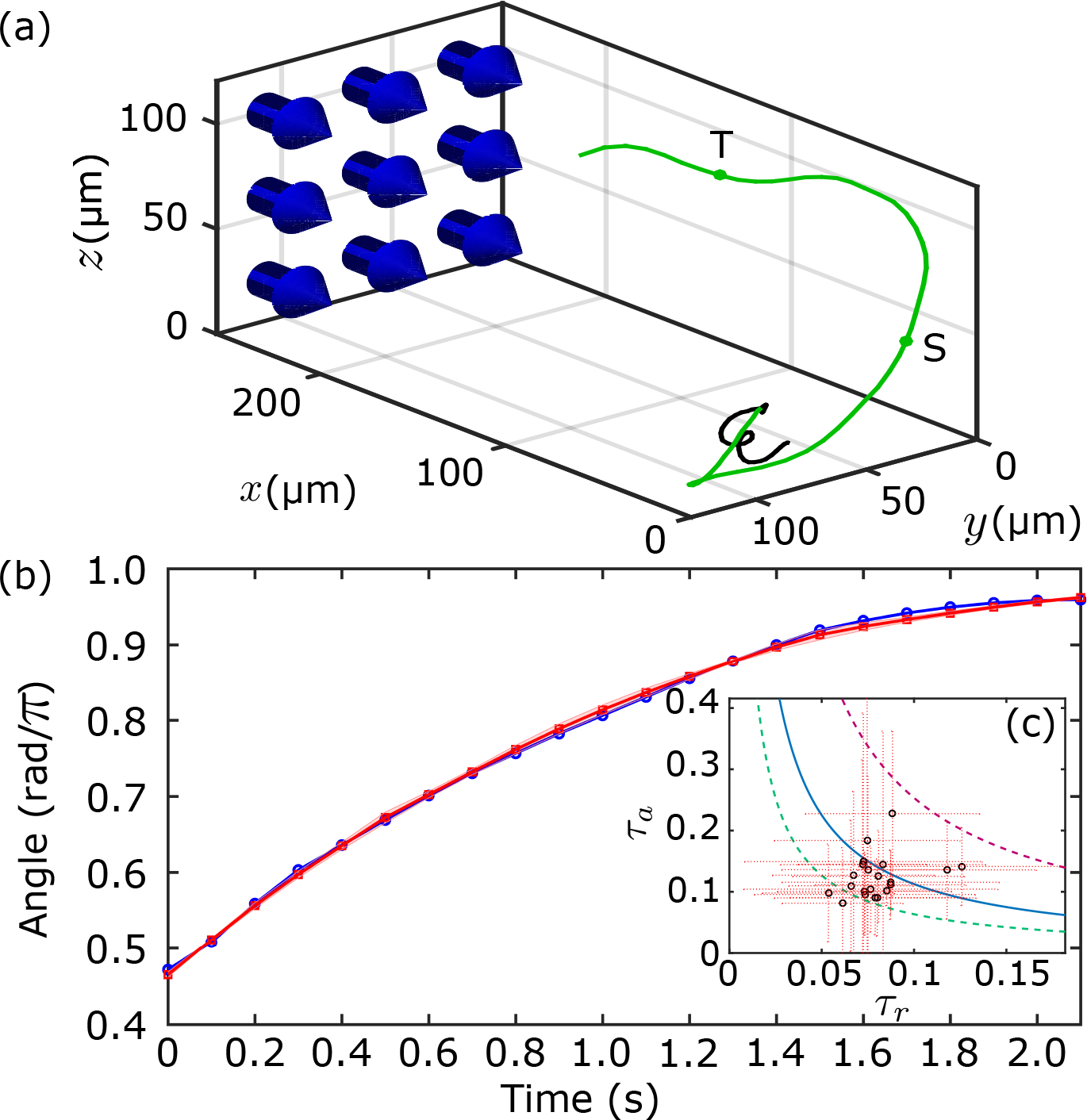
Phototactic swimmers tracked in three-dimensions. (a) The U-turn of a phototactic swimmer shown as acquired in the 3D-tracking apparatus. Trajectory in black indicates the time before light stimulation, whereas trajectory in green indicates after. Blue arrows indicate the direction of the light. The cropped trajectory used for fitting the reorientation dynamics (c) is bounded by the points from S to T. (b) The dynamics of the reorientation angle Ω (in blue) for the cropped trajectory satisfying *π*/2 ⩽ Ω ⩽ *π* (shown in (a) from S to T) fitted by a set of curves (mean and standard deviation in red) described by the iterated map in *Equation 8*, for a convergent range of parameters. (c) Inset showing the means (black markers) of fitted *τ_r_, τ_a_* pairs (standard deviations in red), plotted along the hyperbolas for 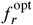 = 1 Hz (red line), 1.5 Hz (blue line) and 2 Hz (green line).

**Figure 6-Figure supplement 1.**
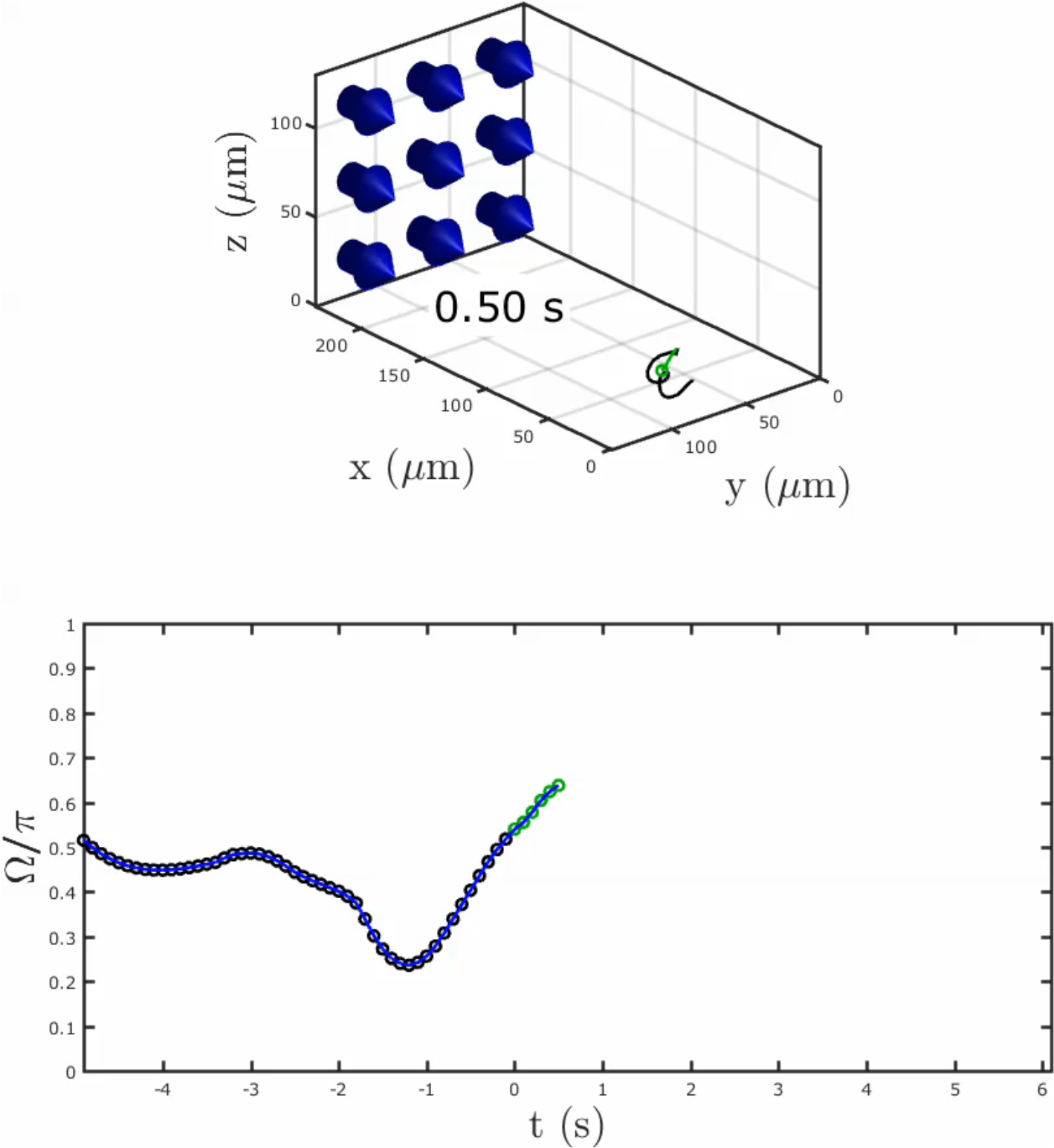
Video of a phototactic swimmer with angle of reorientation. The U-turn of a phototactic swimmer shown in a video with the angle of reorientation plotted below in real time. The colors of the points on the trajectory of the cell before (black) and after (green) the light is on (t = 0) are reflected in the color of the markers on the plot below.

**Figure 6-Figure supplement 2.**
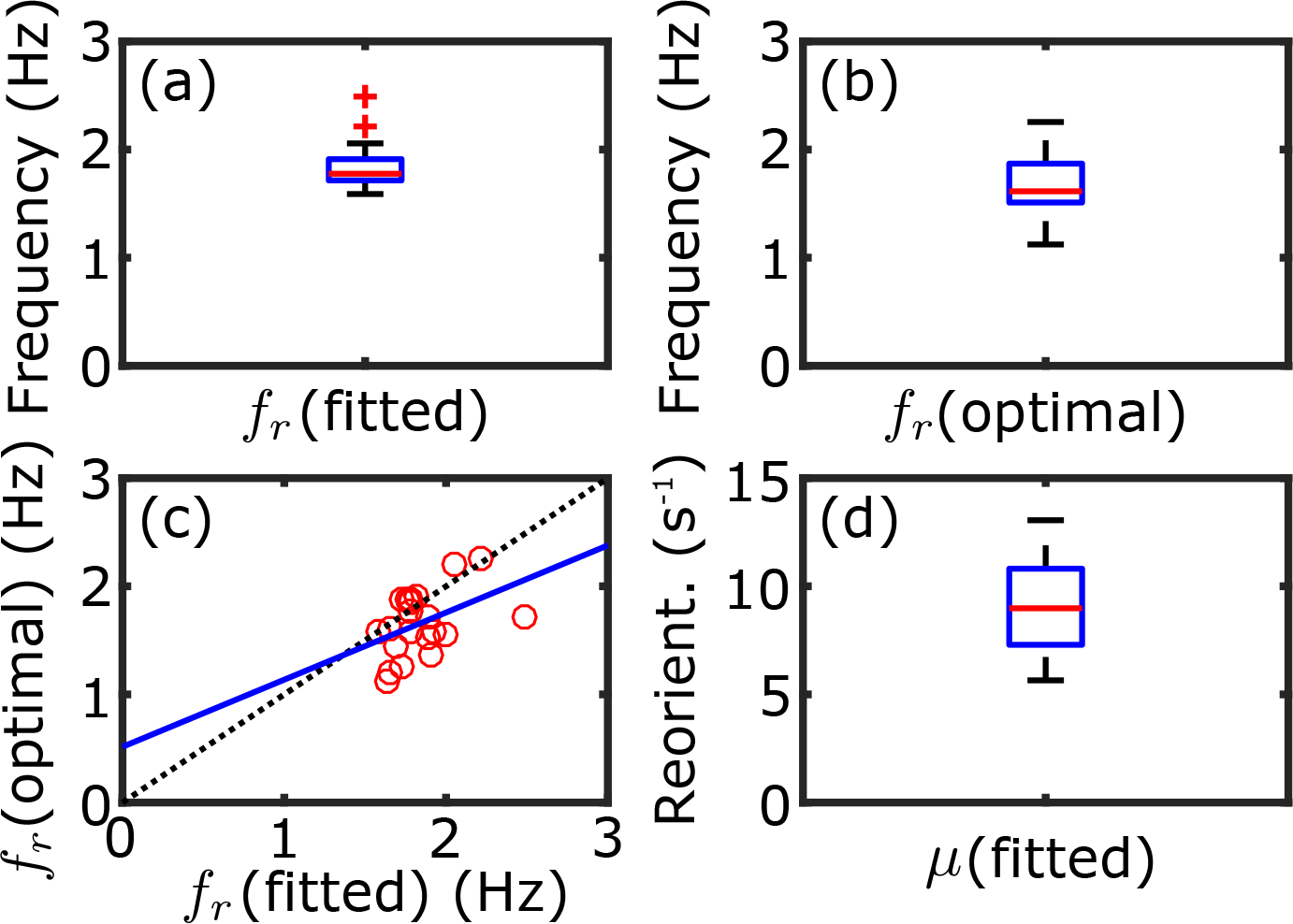
Fitting parameter statistics. (a) Distribution of the fitted rotational frequency with median = 1.78Hz(n = 21). (b) Distribution of the optimal rotational frequency, as defined by **Equation 3** and using the fitted *τ_r_* and *τ_0_* pairs as shown in **Figure 6c,** with median = 1.61 Hz(n = 21). (c) Linear correlation between fitted *f_r_* (from (a)) and optimal *f_r_* (from (b)), shown as a fitted straight line (blue) of the form 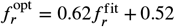. (d) Distribution of the fitted reorientation constant *μ* with median = 8.98 s^-1^ (n = 21).

Regarding the amount of light necessary for a flagellar photoresponse with a positive sign, we have converged, through trial and error, to ≈1 µE·s^-1^·m^-2^ at a wavelength of 470 nm. This value is much lower than in other photoresponse experiments *(Josef et al., 2005)* where ≈60 µE·s^-1^·m^-2^ were used at a longer wavelength (543 nm). This is consistent with the sensitivity profile of channelrhodopsin-2 (*Sineshchekov et al., 2002).* More detailed studies on the wavelength sensitivity of the flagellar photoresponse should be carried out in order to reveal any possible wavelength dependencies on *τ_r_, τ_a_* or *η.*

Our experimental results – coming from different methodologies – show either directly, from the gain of flagellar photoresponse under stimuli of different oscillatory frequencies (*Figure 4a)* or indirectly, from the estimated values of *τ_r_* and *τ_a_* (*Figure 3d, Figure 6c* and *Figure 6-Figure Supplement 2ab)*, that cells with rotational frequency in the range of ≈1-2 Hz would have the most optimal response.

The optimality of the sensitivity of the photoresponse was first addressed by *Yoshimura and Kamiya (2001)*, using a paralyzed-flagella mutant strain *(pf14)* and an electrophysiological approach on a bulk sample. In their experiments, a suspension of immotile cells was exposed to an oscillatlng light stimulus (500 nm) and the resulting photoreceptor current was measured in a cuvette attached to two platinum electrodes. The experiment using relatively high light intensities observed a frequency response peak of 1.6 Hz when stimulated with ≈160 µE·s^-1^·m^-2^ and a frequency response peak of 3.1 Hz when stimulated with ≈40 µE·s^-1^·m^-2^. The former observation is in perfect agreement with our results in *Figure 4a* and in *Figure 6-Figure Supplement 2ab* even though we used light stimulus intensities of ≈1 µE·s^-1^·m^-2^ and ≈5-10 µE·s^-1^·m^-2^respectively. We have not seen any evidence of cells having flagellar photoresponse dynamics that would corroborate the latter result of 3.1 Hz and this is a matter open to further investigation.

Further studies on the optimality of the sensitivity of the photoresponse at the flagellar level were first carried out by *Josef et al. (2006)* on single cells of a negatively phototactic strain. The usage of the quadrature photodiode to measure stroke velocity was vital to the automation of the methodology, nevertheless it gave the magnitude of the velocity component parallel to the body axis only and at a particular position. In this study, it is the first time that the optimality of the photoresponse’s sensitivity is shown in a wild-type strain performing positive phototaxis, both at the flagellar level and at high spatio-temporal resolution, digitally capturing the full waveform of the response.

Moreover, this study addressed the relationship of stimulus *s* to the photoresponse of *Chlamydomonas p* using differential equations and a handful of parameters such as *τ_r_* and *τ_a_* corresponding to physical processes. Attempts to derive similar relationships between stimulus and photoresponse *(Josef et al., 2006)* used linear system analysis. The result of such a signal-processing oriented method, usually includes a much larger number of estimated parameters necessary for the description of the system – without necessarily corresponding to any obvious physical quantities that can be easily measured.

With respect to the range of values observed for *τ_a_* and *τ_r_*, they lie in the low-*τ_r_*/high-*τ_a_* region for step-up, mid-*τ_r_*/mid-*τ_a_* region for 3D-tracking and high-*τ_r_*/low-*τ_a_* region for rotational frequency response experiment. Possible explanations for these observations have to do with the dependence on the intensity of the stimulus (blue) light as well as the interference from the intensity of the background (red) light. It is worthy of commenting that the amount of background light in the immobilized high-resolution experiments is many orders of magnitude higher than the 3D-tracking experiments, and it could very well play a role to the above observations.

The development of a comprehensive mathematical model linking physiology to behavior presents a platform begging for future perturbation-based experiments in order to dissect the mechanism of phototaxis and extend our biological knowledge of the system. The implementation of such a detailed model will require the discovery of many more currently unknown relations between variables, not just for the sake of completeness, but for exploring emerging mechanisms of physiological importance. One such an example is the physiological importance of the parameter of proportionality (*η*) between *p* and *s* (*Equation 1a)* as a measure of phototactic efficiency and phototactic sign, and its dependence on the intensity of the light stimulus.

Flagellar photoresponse – and by extension phototaxis – appears to be a very complex biological process encompassing many variables, as mentioned above. This is evident from the fact that experiments exhibited a high level of difficulty regarding multiple measurements on the same cells of elicited positive photoresponse. This has to do with our lack of understanding of long-term adaptation to darkness or phototactic light for that matter, topics that only recently have begun to be addressed (*Arrieta et al., 2017).*

It is noteworthy to remark that a biochemistry-based model can explain the experimentally observed dynamics of phototactic reorientation in three-dimensions, in the absence of an explicit hydrodynamic model, and with *ω_1_* = -(*1/ζ)p* being sufficient. Although it is evident that the torque generated by each flagellum is connected to the total swept area (*A*) as In *Equation 16* (*Appendix 1)* or to the front amplitudes *(F)* as In the experiments, and that successive differences In the corresponding flagellar torques are responsible for *ω_1_* a more detailed model where biochemistry is coupled to mechanical forces would be the subject of a further study. One example of improving the model could be the investigation of the dependence of flagellar torque to the flagellar beat frequency *(f_b_)* as shown in *Equation 16* (*Appendix 1).* We know from experiments that the frequency of flagellar beat does not change significantly during the photoresponse experiments on immobilized cells (*Figure 3-Figure Supplement 3* showing the cell with the most change to be ⪅ 10%), but not necessarily for free-swimming cells. Another example of including more detailed hydrodynamics, would be the formal definition of *1/ζ*, the proportionality constant between *ω_1_* and *p.* Interestingly from the fitted parameter *μ*(= *σ/ζ* in *Equation 21)*, we know that the product between *σ* and *1/ζ* is of order 10 (median value 8.98 s^-1^) and although we do not know the exact value of *σ* (= *s_0_η)*, we can estimate it to be in the range of 4 < *σ <* 7 based on the light intensities used. This allows us to place an estimate on 1/*ζ* in the range of 1.3 < 1/*ζ* < 2.3, and if we compare it to the relation *ω_1_ = –(τ/ζ_r_)(2b/a)* we can further relate It to the fluid mechanics via 1/*ζ* = τ*/ζ_r_* × *F/A* = 5.7 s^-1^, where *F/A* is empirically found to be ≈1/3.5. In this study, we declare this level of proximity, i.e. same order of magnitude, between observed and expected values of *1/ζ*, a success, and we leave a more accurate estimate of this variable to future, more detailed hydrodynamic models that similarly link physiology to behavior.

## Methods and Materials

This is a detailed description of the materials and methods used for both types of experiments with immobilized and free-swimming cells and their corresponding analyses.

### Culture conditions

*Chlamydomonas* wild-type cells (strain CC125 (*Harris, 2009))* were grown axenically under photoautotrophic conditions in minimal media (*Rochaix et al., 1988)*, at 23°C under a 100 μE·s^-1^·m^-2^ illumination in a 14:10 h light-dark cycle.

### Flagellar photoresponse of immobilized cells

Cells were prepared as described previously (*Leptos et al., 2013)* – centrifuged, washed and gently-pipetted into a custom-made observation chamber made of polydimethylsiloxane (PDMS) as shown In *Figure 2a.* Chambers were mounted on a Nikon TE2000-U inverted microscope with a x63 Plan-Apochromat water-immersion long-working-distance (LWD) objective lens (441470-9900; Carl Zeiss AG, Germany). Cells were immobilized via aspiration using a micropipette (B100-75-15; Sutter, USA) that was pulled to a ø5-µm tip, and the flagellar beating plane was aligned with the focal plane of the objective lens via a rotation-stage. Video microscopy of immobilized cells was performed using a high-speed camera (Phantom v341; Vision Research, USA) by acquiring 15 s-movies at 2000 fps. Cells were stimulated at exactly frame 2896 (≈1.45 s into the recording) using a ø*50* µm-core optical fiber (FG050LGA; Thorlabs, USA) that was coupled to a 470 nm Light Emitting Diode (LED) (M470L3; Thorlabs, USA) and was controlled via an LED driver (LEDD1B; Thorlabs, USA). The LED driver and the high-speed camera were triggered through a data-acquisition card (NI PCIe-6343; National Instruments, USA) using in-house programs written in `LabVlEW 2013` (National Instruments, USA), for both step- and frequency-response experiments. Calibration of the optical fiber was performed as follows: A photodiode (DET110; Thorlabs, USA) was used to measure the total radiant power *W* emerging from the end of the optical fiber for a range of voltage output values (0-5 V) of the LED driver. Subsequently, the two quantities were plotted and fitted to a power-law model which was close to linear.

A stimulus of ≈1 µE·s^−1^·m^−2^(at 470 nm) was empirically found to give the best results in terms of reproducibility, sign, i.e. positive phototaxis, and quality of response, since we conjecture that the cells could recover in time for the next round of stimulation. For the step response experiments, biological replicates were *n*_cells_ = 4 with corresponding technical replicates *n*_tech_ = {4,3,2,2}. For the frequency response experiments, biological replicates were *n*_cells_ = 3 with each cell stimulated to the following amplitude-varying frequencies: 0.5 Hz, 1 Hz, 2 Hz, 4 Hz and 8 Hz. Only the cells that showed a positive sign of response for *all* 5 frequencies are presented here. This was hence the most challenging aspect of the experimental process.

### Analysis of flagellar photoresponse

High-speed movies were processed and flagellar features were extracted as described previously (*Leptos et al., 2013).* The angle Θ (*Figure 3-Figure Supplement 1*) between a flagellum chord (i.e. the line connecting the base of the flagellum and a point at a fixed distance from the base) and the axis of symmetry of the cell was used to define the duration of the flagellar beats. In particular, the beginning and the end of the beat were defined by the local minima in a time-series of the angle Θ *(Figure 3-Figure Supplement 1).* For every *in-phase* beat, the areas swept by the two flagella and located above the two reference lines drawn at 60° from the cell’s central axis (noted as F_cis_ and F_trans_) were measured. These are shown in *Figure 3a-b*, and were used as the front amplitudes for each beat. Finally, the flagellar photoreponse was defined as the difference of normalized front amplitudes, where the normalization factor was the average front amplitude for the corresponding unstimulated cell. The front amplitude (F) of beats during instantaneous asynchronies were ignored and the corresponding values at those points were interpolated.

The solution to the governing equations (*Equation 1a* and *Equation 1b)* for an oscillatory stimulus with frequency *f_s_(= ω_s_/2π)* such as

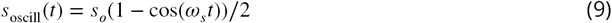

can be written in closed form (for sufficiently large enough *t*):

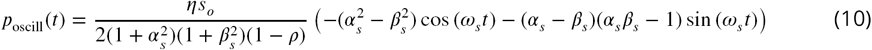

where

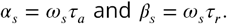

The gain of frequency response is thus defined as the magnitude ratio

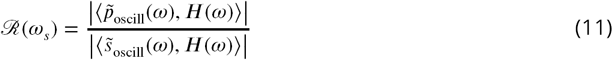

where 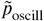 and 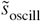 are the Fourier transforms of *p*_oscill_ and, *s*_oscill_ respectively. Truncation for positive frequencies is indicated by 〈*·,H(w)〉.*

### Phototaxis experiments of free-swimming cells

Three-dimensional tracking of phototactic cells was performed using the method described in *Drescher et al. (2009)* and shown in *Figure 2b.* The experimental setup comprised of a sample chamber suspended in an outer water tank to eliminate thermal convection. The sample chamber was composed of two acrylic flanges (machined in-house) that were clamped onto an open-ended square borosilicate glass tube (2cm × 2 cm × 2.5 cm; Vetrospec Ltd, UK), in a watertight fashion. Two charge-coupled device (CCD) cameras (Prosilica GT750; Allied Vision Technologies, Germany) coupled with two InfiniProbe™ TS-160(Infinity, USA) with Micro HM objectives at a total magnification of x16. The source of phototactic stimulus was a 470 nm blue-light LED (M470F1; Thorlabs, USA) coupled to a solarization-resistant optical fiber (M22L01; Thorlabs, USA) attached to an in-house assembled fiber collimator that included a ø12.7 mm plano-convex lens (LA1074-A; Thorlabs, USA). Calibration of the collimated optical fiber was performed similarly to the experiments with immobilized cells. In addition, the thickness of the walls of the outer water tank, the walls of the inner sample chamber and the water in between, were taken into account for the calibration.

The two CCD cameras and the blue-light LED used for the stimulus light were controlled using `LabVIEW 2013` (National Instruments, USA) including the image acquisition driver `NI-IMAQ` (National Instruments, USA). The cameras were triggered and synchronized at a frame rate of 10 Hz via a data-acquisition device (NI USB 6212-BNC; National Instruments, USA). For every tracking experiment (M = 6), two 300-frame movies were acquired (side and top) with the phototactic light triggered at frame 50 (5 s into the recording). The intensity of the blue-light stimulus was chosen to be 5 or 10 µE·S^−1^·m^−2^.

### Analysis of three-dimensional tracks

To track the cells we used in-house tracking computer programs written in **MATLAB** as described in *Drescher et al. (2009).* Briefly, for every pair of movies cells were tracked in the *side* and *top* movies corresponding to the *xz*-plane and in the *xy*-plane respectively. The two tracks were aligned based on their *x*-component to reconstruct the three-dimensional trajectories. The angle Ω *(Figure 6b)* between the cell’s directional vector and the light was then calculated for every time point. The post-stimulus sections of the trajectories were cropped to the interval *π*/2 ⩽ Ω ⩽ *π*, which corresponds to the reorientation phase. Using the relation Ω = Φ + *π/2*, the cropped trajectories were fitted to *Equation 8* by estimating the following parameters *τ_r_, τ_a_, f_r_* = *ω_r_*/2*π* and μ = *σ/ζ.* The deterministic Nelder-Mead simplex method was employed to minimize the residual sum of squares *(RSS).* In order to avoid parameter estimations associated with local minima, 3000 different initial-condition vectors of the form (*τ_r_, τ_a_, f_r_, μ*)^init^ were used for the fitting of each trajectory. These vectors were constructed using all possible permutations from the following sets: 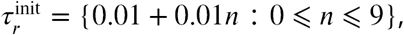 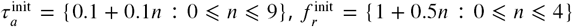 and 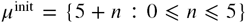 such that 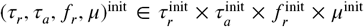. The criterion of good fit was taken to be RSS *<* 0.03 for at least 5% (i.e. 600) of the fitting attempts using different initial conditions.

## Acknowledgments

We would like to thank Pierre A. Haas for very useful discussions regarding mathematical theory, advice on Euclidean geometry, and critical reading of the manuscript, Kirsty Y. Wan for sharing some initial code from previous work on flagellar tracking, David-Page Croft, Colin Hitch and Paul Mitton in the mechanical workshop at DAMTP for technical support, John Milton for support with electronics, also at DAMTP, and Ali Ghareeb for helping with the initial assembly of the fiber coupling apparatus.

## APPENDIX 1 Calculations used in estimating the angle of phototactic turning Derivation of time-averaged total torque generated by rod-shaped flagella

In order to derive the amount of phototactic turning per half turn of cell rotation, we consider a swimmer in Stokes flow with a spherical body of radius *R*, bearing two rod-shaped flagella of length *L* attached at the anterior of the cell body, as shown in *Appendix 1-Figure 1*. The swimmer is immersed in a fluid with viscosity *η*. Furthermore, the swimmer flaps its rod-shaped flagella with a maximum velocity at the tip equal to

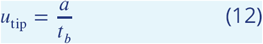

where *a* is the amplitude of the beat and *t_b_* is the duration of the effective stroke of the beat. We can thus assign each flagellum a force-density function depending on the position *λ* along the flagellum:

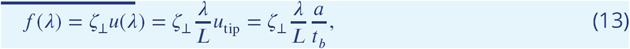

**Appendix 1 Figure 1.**
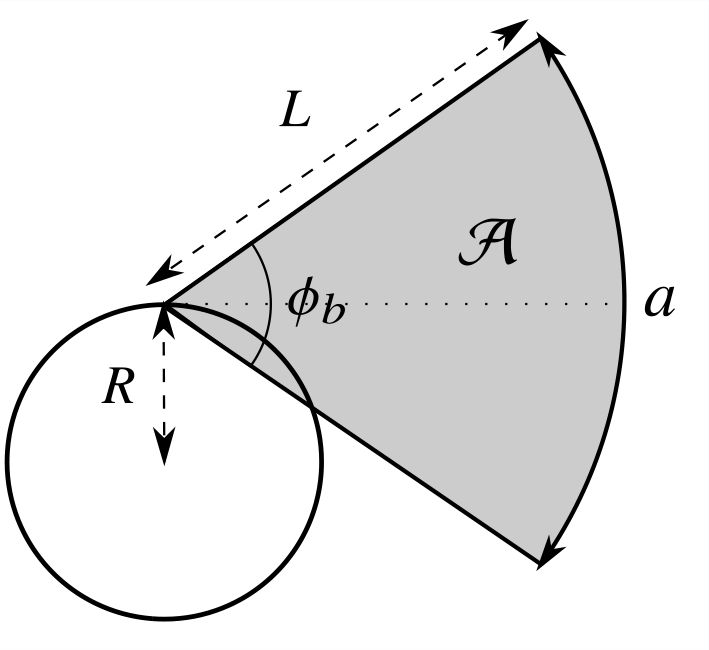
Model of the effective stroke of a simplified swimmer. The total angle spanned by the rod-shaped flagella during an effective stroke is equal to *ϕ_b_* = *a/L* and the corresponding swept area (shaded) is 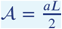

**Appendix 1 Figure 2.**
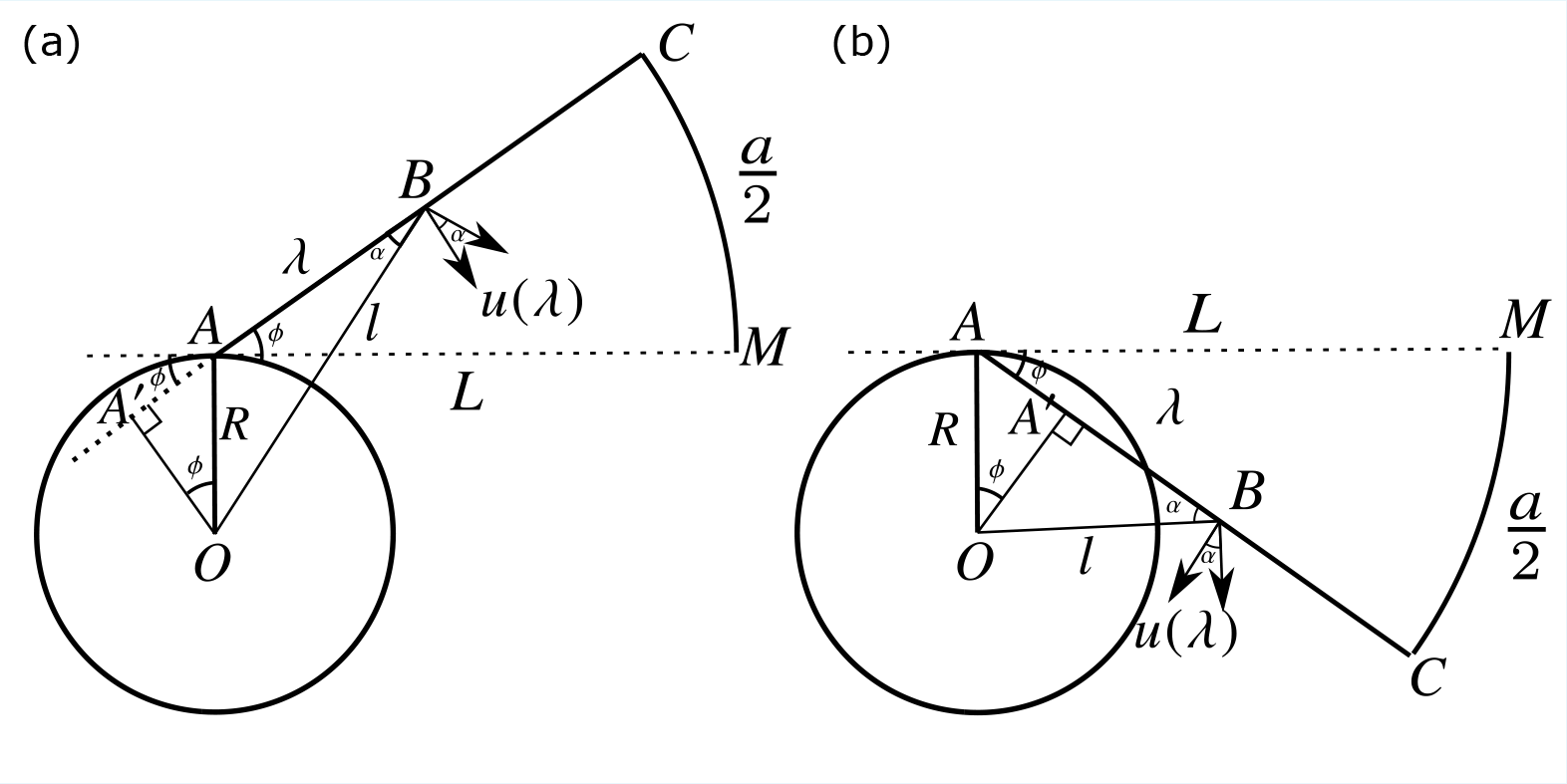
Geometric proof of the torque generated by a point on the rod-shaped flagellum. The proof is broken down in two parts: (a) for angles of *ϕ* above the tangent *AM* to the cell body (front amplitude) and (b) for angles of *ϕ* below the same line (back amplitude).

The corresponding torque density, according to the geometric proof shown in *Appendix 1-Figure 2a*, for the front amplitude is

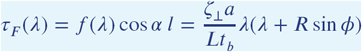

where *OA = R, AC = AM = L, AB = λ, OB = l and l cos α = BA’ = AB + AA’ = λ + R* sin *ϕ*.

The corresponding torque density, according to the geometric proof shown in *Appendix 1-Figure 2b*, for the back amplitude is

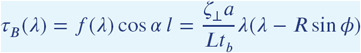

where *OA = R, AC = AM = L, AB = λ, OB* = l and *l cos α = BA’ = AB – AA’ = λ – R* sin *ϕ.*

The torque density functions are also functions of *t*, as *ϕ* is a function of *t.* We define 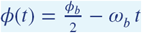 for 0 ⩽ *t* ⩽ *t_b_*, where 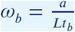. Thus *τ_F_* and *τ_B_* can be combined and rewritten as

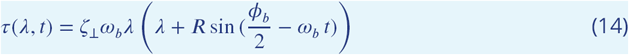

Then the time-averaged total torque generated by a flagellum during the effective stroke of the beat is equal to

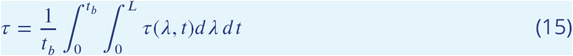

Performing the computation yields

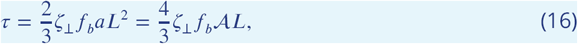

where *t_b_* = 1/*f_b_* and *f_b_* is the frequency of beating, and *A = aL/2* is the area of the circular sector swept by the flagellum.

### Estimate of perpendicular drag coefficient

Using the definition of ζ_⊥_ described in *Pak et al. (2011)*

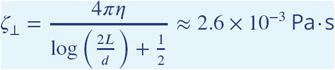

using *d =* 0.25 µm for the diameter of the flagellum and *η* = 10^−3^ Pa·s as the viscosity of the fluid.

### Estimate of rotational drag coefficient

Using the values above for *a* and *η* we calculate *ζ_r_* to be

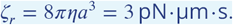

### Integration of oscillating angular velocity

We would like to estimate the angle by which the cell turns – about its 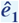 axis – for the duration of half a turn about its 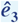 axis, while its angular velocity *ω_1_* oscillates as shown in *Equation 6.* It is reasonable to assume that the difference In flagellar amplitude between the two flagella *(2b)*, on which *ω_1_* depends, oscillates with constant amplitude (2*b_0_) during the period of half a turn.* Then we can compute the angle turned about 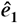 during the time the cell turns by an angle *π* about 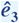 by integrating *Equation 6* over time from to 1/2*f_r_:*

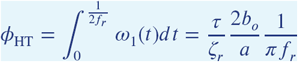

If we substitute for *τ* then we have

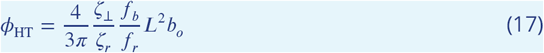

## APPENDIX 2 Derivation of mathematical model

The mathematical model is derived from a system of five nonlinear ordinary differential equations (ODEs) following a series of simplifications and approximations. The first simplification is regarding the photoresponse time delay *t_d_*, as mentioned in the main text. We know from solving the equations numerically that including the time delay into the mathematical model is equivalent to omitting it, but with the eyespot vector *ô* lying on the 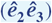 plane, i.e. 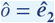 (*Figure 1a).*

More specifically, the dynamics of the photoresponse (described by *Equation 1a* and *Equation 1b)* are coupled to the Euler angle dynamics via the light intensity relation 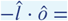 = –*s_o_* (sin *ψ* cos *ϕ* + cos *θ* sin *ϕ* cos *ψ*), where 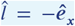 (*Figure 6a)*, and the equations describing the Euler angle dynamics (*Symon, 1971)* are coupled to the photoresponse via the relation *ω_1_* = -(1/*ζ)p*, where *ζ* is a time scale constant equal to an effective viscosity. This gives the following system of ODEs:

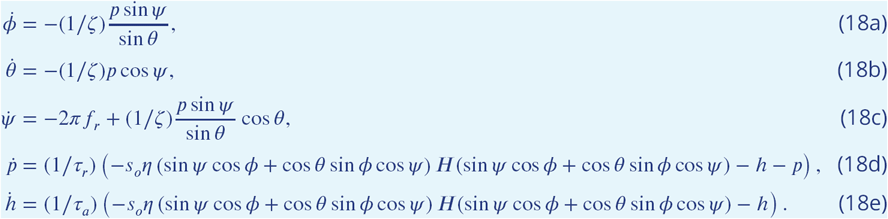

Using the test case where the initial direction of the cell is 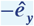 i.e. Euler angle initial conditions *θ* = *π/2* and *ϕ =* 0, we conclude from the solution of the reorientation dynamics that the cell maintains a trajectory on the (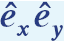) plane with *θ* being almost constant. With *θ* being constant and *ψ* = -2*πf_r_t (Figure 1)* we can reduce the number of equations in the system from five to three. Additionally with the nondimensionalization of time 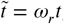, where *ω_r_ = 2πf_r_* the equations transform to

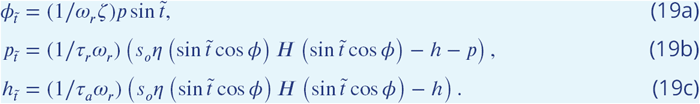

To be able to decouple *ϕ* from *h* and *p*, we assume that it does not change significantly during a full (or half) cell rotation, and thus we solve the equations for *h* and *p* for a given value of *ϕ.*

If we let *α = τ_a_ω_r_, β* = *τ_r_ω_r_*, σ = *s_0_η* and dropping tildes, we can rewrite the equations as follows:

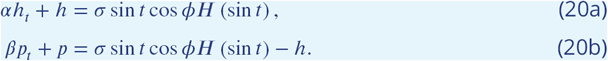

***Equation 20a*** can be rewritten as,

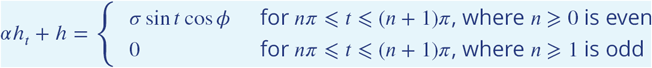

and it can be solved in a piecewise fashion to yield,

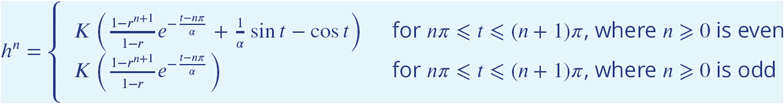

where

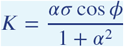

and

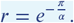

Likewise, *Equation 20b* can be rewritten more analytically as

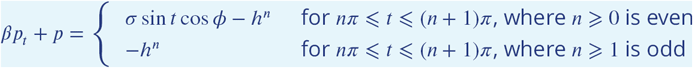

and it can be solved in a piecewise fashion as to yield.

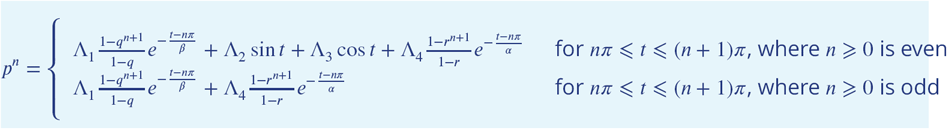

where

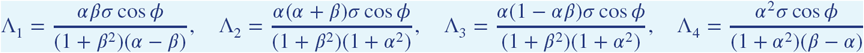

and

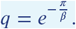

Since *n* represents the number of half-turns, where for even values it corresponds to the times where the cell’s eyespot is receiving light and for odd values to the times where the cell is in the “darkness”, we integrate *Equation 19a* for every value of *n* ⩾ 0

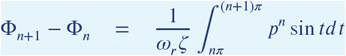

which can be written in the form of *Equation 8*, where

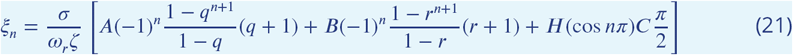

and where

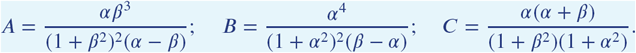

